# A biophysical framework to accurately identify antigen single-amino acid escape variants and corresponding variant-specific compensatory TCR sequences

**DOI:** 10.64898/2026.02.02.703411

**Authors:** Zahra S. Ghoreyshi, Herbert Levine, Xingcheng Lin, José N. Onuchic, Jason T. George

**Affiliations:** Department of Biomedical Engineering, Texas A&M University, College Station, TX, USA; Translational Medical Sciences, Texas A&M University Science Center, Houston, TX, USA; Center for Theoretical Biological Physics, Rice University, Houston, TX, USA; Center for Theoretical Biological Physics, Northeastern University, Boston, MA, USA; Department of Physics; Department of Bioengineering, Northeastern University, Boston, MA, USA; Department of Physics, North Carolina State University, Raleigh, NC, USA; Bioinformatics Research Center, North Carolina State University, Raleigh, NC, USA; Department of Physics and Astronomy, Department of Biosciences, and Department of Chemistry, Rice University, Houston, TX, USA; Department of Hematopoietic Biology and Malignancy, UT MD Anderson Cancer Center, Houston, TX, USA

## Abstract

The impact of single amino acid substitution on T-cell receptor (TCR) recognition is central to understanding the molecular determinants of TCR specificity and degeneracy during viral mutational escape, cancer recognition, and autoimmunity. In this study, we developed a biophysics-informed computational approach integrating experimental alanine-scan mutagenesis data from the autoimmune-associated ALWGPDPAAA peptide bound to HLA-A*02:01 together with coarse-grained structural modeling. Our approach reconstructs the energetics and structural determinants underpinning the observed loss of recognition by the diabetogenic 1E6 TCR upon single-point mutations, specifically at the critical Pro^5^ and Asp^6^ residues. Leveraging the computational model’s ability to incorporate multiple structural templates into binding predictions, this approach quantitatively reproduces experimentally measured affinity disruptions. Additionally, we apply our approach to identify potential compensatory interactions capable of restoring binding affinity through alternative residue interactions. Application of our framework to independent CMV and SARS-CoV-2 peptide-specific TCRs yielded accurate specificity predictions on the associated peptide point mutants. This integrative computational framework contributes a strategy for inferring TCR-peptide binding energetics at the single amino acid level, guiding the rational design of peptide-based immunotherapeutics, and predicting the functional impacts of clinically relevant peptide variants.

## Introduction

Understanding the molecular interactions underlying T-cell receptor (TCR) recognition of peptide-major histocompatibility complex (pMHC) complexes is essential for elucidating fundamental principles of immune specificity and tolerance, as well as for informing the design of novel immunotherapeutic strategies [1]. In this context, detailed knowledge of the structural and energetic determinants governing TCR specificity against pMHC can provide critical insights into various disease processes, including viral and tumor recognition, autoimmunity, and host-donor compatibility.

Single–amino-acid peptide variants (SAPVs) dominate many clinically relevant T-cell epitopes. In viruses, a single residue change can abolish recognition by previously protective cytotoxic T cells and drive immune escape, as documented for HIV and SARS-CoV-2 [2]. In oncology, tumor neoantigens arise from non-synonymous point mutations, providing the basis for personalized cancer vaccines and checkpoint-blockade responses [3]. In transplantation, minor histocompatibility antigens most often arising from single nucleotide polymorphisms (SNPs) elicit potent alloreactive T-cell responses that influence graft-versus-host disease risk [4]. Reliable prediction of TCR recognition across this vast SAPV landscape is therefore central to viral-variant surveillance, neoantigen prioritization, and donor–recipient matching in transplantation.

A growing body of computational work has sought to model and predict TCR–pMHC recognition, spanning purely data-driven, structure-based, and hybrid approaches. Sequence-focused neural network models such as TULIP [5], DeepTCR [6], ImRex [7], and BATMAN [8] learn TCR specificity directly from large repertoire datasets without explicit structural information, enabling high-throughput prediction but often lacking physical interpretability. Structure-oriented frameworks, including STAPLER [9] and template-based docking or geometric alignment models [10], estimate binding geometry and interface complementarity using structural templates. Hybrid energy-guided approaches— including RACER [11], and related biophysical extensions [12–16]— combine coarse-grained energetics with structural features or machine learning to capture both the geometry and physics of TCR recognition. Collectively, these methods have substantially advanced our ability to decode TCR specificity across viral, tumor, and autoimmune contexts. Building on this foundation, RACER-m [17] introduces a compact, physics-informed model that leverages structural energetics to predict not only binding affinity but also compensatory mechanisms of antigenic escape, thereby bridging the gap between statistical prediction and mechanistic understanding.

Despite these advances, deep learning approaches trained predominantly on bulk TCR–pMHC binding data, including the recent repertoire-scale classifiers and embedding-based architectures discussed above, remain limited in their ability to resolve the fine-grained energetic consequences of single–amino–acid substitutions. Such models are highly effective for broad antigen classification, repertoire clustering, and large-scale specificity inference, yet they are not optimized to capture point-mutation effects that critically alter recognition at individual peptide positions, an aspect central to viral escape and tumor neoantigen recognition, where a single residue change can toggle TCR activity. In contrast, physics-informed, structure-based energy models such as RACER-m [17] directly encode residue-level interactions and local interfacial geometry, providing interpretability and sensitivity to the subtle structural perturbations induced by SAPVs. As a result, these mechanistic models offer a complementary and mutation-aware alternative to purely data-driven deep learning frameworks.

To address the limitations of current approaches, we developed a biophysics-informed computational pipeline leveraging experimental alanine-scan mutagenesis data from the autoimmune-associated ALW peptide bound to HLA-A*02:01, combined with the coarse-grained structural modeling capabilities of RACER-m [17]. Our integrative approach reconstructs the structural and energetic landscape underpinning the experimentally observed loss of 1E6 TCR recognition with mutations of the critical Pro^5^ and Asp^6^ residues [18]. By utilizing RACER-m’s multi-template threading and residue-level energetic calculations, our model quantitatively reproduces experimentally measured disruptions in TCR-pMHC affinity. Furthermore, this approach enables the identification of compensatory interactions that could potentially restore affinity, providing insight into alternative molecular strategies to modulate auto-reactive TCR responses.

Collectively, this integrative computational framework advances our understanding of TCR-pMHC recognition, offering robust methodologies for dissecting the energetics of immune receptor interactions. This approach has significant implications for guiding the rational design of peptide-based immunotherapy and for predicting the functional consequences of clinically relevant peptide variants. Finally, we confirm the generality of our methods by showing that the approach developed here can lead to successful predictions for two other cases of TCRs tested against a range of mutated antigens.

## Methods

### Computational modeling of TCR-pMHC interactions using energy-based model

To systematically explore the effects of point mutations on TCR recognition of pMHC complexes, we employed a computational framework based on the RACER-m energy model [17]. This approach calculates residue-level interaction energies between TCR and peptide primary amino acid sequences, which can be computed as:

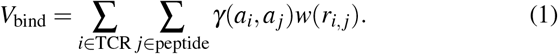

Here, *a*_*i*_ and *a* _*j*_ are the amino acid types at TCR residue *i* and peptide residue *j, γ*(*·, ·*) defines the residue-residue interaction potential between TCR and peptide amino acids, and *r*_*i, j*_ is the inter-residue distance (*C*_*β*_ -*C*_*β*_ ; *C*_*α*_ for glycine), and *w*(*r*) is a smooth and decreasing distance-based weighting based on amino acid proximity.

Here, *γ* is a single 20*×* 20 matrix of residue–residue interaction coefficients, one entry per amino-acid pair, shared across all TCR–peptide residue pairs in Eq. (1); it therefore depends only on which two amino acid types are in contact, not on sequence position. Following RACER-m [17], the entries of *γ* are learned by an inverse-statistical (energy-based) optimization procedure that adjusts the matrix so that experimentally known strong-binding TCR–pMHC complexes in the training set score more favorably than a large ensemble of decoy (peptide-randomized) complexes for the same TCR; the same trained *γ* matrix is then reused, unchanged, to score every peptide and CDR3 mutant considered in this study. Retraining *γ* (as in the antigen-restricted and AlphaFold3-augmented configurations described below) means re-running this same optimization procedure on an expanded or restricted set of training complexes, rather than changing the functional form of Eq. (1) itself.

### Structure preparation and peptide mutation

For each wildtype (WT) TCR–pMHC complex, we enumerated all single–amino acid substitutions across the peptide by replacing the native residue at each position *k* with each of the 19 non-native amino acids. To construct 3D models for every (TCR, peptide^mut^) pair, we adopted RACER-m’s template-based threading strategy [17]. Briefly, among a reference set of solved TCR–pMHC structures, we computed position-wise Hamming similarity between the target sequences and each structural template for the peptide, CDR3*α*, and CDR3*β* segments. We then formed a composite similarity score

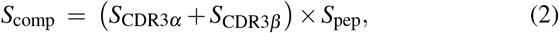

and selected the structural template that maximized *S*_comp_ for downstream modeling. Following RACER-m, threading was performed in MODELLER by replacing the template peptide, CDR3*α*, and CDR3*β* sequences with the target sequences, aligning the first residue and padding any length mismatch with gaps before model building; the resulting structure was used for energy evaluation.

### Energy minimization and refinement

For each threaded TCR–pMHC complex, the predicted binding energy is evaluated using the coarse-grained pairwise interface potential defined in Eq. 1. The following sigmoidal window was implemented:

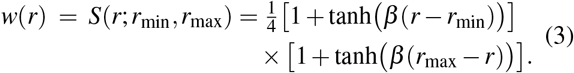

with *r*_min_ = 6.5 Å, *r*_max_ = 8.5 Å, and slope *β* = 5 Å^*−*1^. Only peptide residues and TCR interface residues (primarily CDR3*α/β* regions) are included in the sums of Eq. (1). Because evaluation of the interface potential requires a complete TCR–pMHC structural model, the *In silico structure generation* section describes how such complexes are generated *in silico* when peptide mutations necessitate new geometries.

### In silico structure generation

To determine when additional antigen-specific structures were required, we assessed the structural coverage provided by the available TCR–pMHC complexes together with the model’s ability to reproduce the experimentally observed mutational landscape. When additional structural coverage was needed, representative experimentally supported strong-binding peptide variants were selected for structure generation and subsequently incorporated into the antigen-restricted training set. The overall decision and structural-augmentation workflow is summarized in Supplementary Figure S8.

For analyses requiring peptide-side structural variation, full TCR–pMHC complexes were generated *in silico* using AlphaFold3. For each peptide sequence of interest, complete variable-region sequences for TCR*α* and TCR*β* were assembled by retaining the V*α* and V*β* framework regions from the crystallographic 1E6 structure (PDB: 3UTT) and substituting only the CDR3 loops and peptide segment according to the specified mutation. The HLA-A*02:01 heavy chain was kept fixed across all models to preserve a consistent MHC framework.

Each assembled TCR–pMHC sequence was submitted to the AlphaFold3 pipeline using standard inference settings. Predicted structures were examined to confirm backbone continuity and to verify that the overall TCR–peptide–MHC geometry remained compatible with the established 1E6 docking orientation. The resulting *in silico* complexes were then passed directly into the RACER-m refinement and scoring workflow described above. When additional experimentally determined or AlphaFold3-derived structures were incorporated into training, RACER-m was updated by reoptimizing the same pairwise residue–residue interaction matrix using the expanded set of complexes, while keeping the functional form of the coarse-grained potential unchanged.

We use two distinct structure-generation strategies at different stages of the pipeline for practical reasons rather than as independent validations of one another. Template-based threading (*Structure preparation and peptide mutation*, above) is computationally lightweight and is therefore used for the exhaustive, combinatorial enumeration of single-point peptide substitutions and, separately, single-site CDR3 substitutions; this scale is not currently practical with AlphaFold3. AlphaFold3 modeling, in contrast, is applied only to a small number of representative strong-binding mutants and predicted compensatory complexes, where it serves as an independent, non-threading-based geometric plausibility check on cases already prioritized by RACER-m. The two approaches are compared directly via mutual *Q* analysis between threaded and AlphaFold3-derived structures for the same sequences (Supplementary Figures S3–S5), which we use to assess the degree to which the two structure-generation methods agree.

### Standardized Scoring and Classification Criteria

We note that the experimental readouts used to define “strong” and “weak” binders differ across the TCR–pMHC systems considered in this study. For 1E6–ALW, binding classes are defined using TNF*α* release, whereas for the NLVPMVATV-specific TCR81-14 dataset, the experimental readout is NFAT luminescence. Both TNF*α* release and NFAT luminescence measure functional T-cell activation rather than TCR–pMHC binding affinity directly. In contrast, the SARS-CoV-2 SVAR-73 dataset [19] uses pMHC tetramer binding as the experimental readout, providing a more direct measure of TCR– pMHC recognition. Functional activation depends not only on TCR–pMHC binding energetics, but also on binding kinetics, peptide–MHC stability and presentation, TCR and coreceptor abundance, and downstream signaling thresholds. RACER-m’s energy scale (Eq. (1)) is intended as a proxy for TCR–peptide binding energetics. Accordingly, comparisons with TNF*α* release or NFAT luminescence test whether this energetic proxy tracks functional activation, whereas comparison with the SVAR-73 tetramer-binding data more directly tests whether the predicted energetic landscape tracks experimentally observed TCR–pMHC recognition.

To place energies on a common reference scale, each test TCR–peptide complex is scored against a peptide-randomized *<* decoy *>* ensemble of size 1000 (per complex). Let *µ*_decoy_ and *σ*_decoy_ denote the sample mean and standard deviation of the 1000 decoy energies for that fixed TCR. The standardized score for a test energy *E*_test_ is

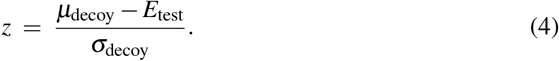

Decision thresholds are calibrated from the WT distribution computed on the 1E6 training templates. Here, *µ*_WT_ denotes the average WT *z*-score across templates in the training set (the WT reference margin), and *σ*_WT_ denotes the standard deviation of those WT *z*-scores (the spread of the margin). We define *z*_crit_ = *µ*_WT_ *− σ*_WT_, which sets the threshold one standard deviation below the WT reference margin. Classification is then

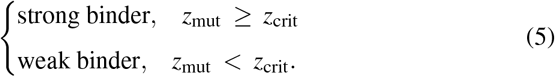

Thus, a peptide point mutant is flagged as disrupted when its *z*-score falls below *z*_crit_, with all quantities defined on the same decoy-normalized scale (Eq. (4)). Because RACER-m models only TCR–peptide energetics, the next section evaluates the contribution of peptide–MHC binding stability to the observed mutational effects.

### Evaluation of peptide–MHC interactions

Because RACER-m explicitly models only TCR–peptide interaction energies, we separately evaluated the contribution of peptide–MHC stability to the ALW mutational landscape. For each single–amino-acid substitution, peptide–MHC binding affinity was estimated using NetMHCpan 4.1 [20], which provides allele-specific predictions for HLA-A*02:01. These values were used to identify peptide variants whose experimental loss of function is more likely attributable to impaired MHC presentation than to altered TCR engagement.

Positions known from structural data to be strongly MHC-facing (Leu^2^, Pro^7^, Ala^9^, Ala^10^) were examined in detail by comparing their predicted stability changes with RACER-m *z*-scores and experimental TNF measurements. This auxiliary analysis enabled a mechanistic separation of TCR-driven versus MHC-driven effects and informed interpretation of mutational outcomes that cannot be captured solely from TCR–peptide energetics.

Throughout this work, we refer to this composite scoring strategy—RACER-m’s coarse-grained TCR–peptide energetics, augmented with AlphaFold3-derived structural modeling at TCR-facing positions and with NetMHCpan-based peptide–MHC affinity prediction at MHC-facing positions—as RACER-MAP.

### Identification of compensatory interactions

To determine whether TCR-side changes can rescue specificity lost after peptide point mutations, we performed an exhaustive single-site scan over the CDR3 loops of both chains. For each peptide mutant previously classified as <u>disrupted</u> (i.e., *z*_mut_ *< Z*_crit_), we generated the one-mutation neighborhood of the TCR, defined as

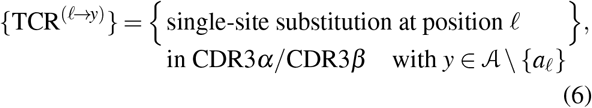

where A denotes the set of 20 standard amino acids and *a*_*l*_ is the WT residue at position *l*. Each candidate TCR variant was then paired with the peptide mutant to form complexes of the form (TCR^(*l→y*)^, *p*^(*k→x*)^).

Each candidate complex was modeled using the same template-selection, threading, and refinement protocol described above: (i) an optimal structural template was selected via the composite sequence-similarity score (peptide, CDR3*α*, CDR3*β*), (ii) mutated CDR3 segments and peptide were threaded with MODELLER, preserving backbone geometry, and (iii) an identical minimization and interface-energy evaluation (including decoy normalization with 1000 peptide-randomized decoys per fixed TCR) was applied, as performed for both the WT and peptide mutants. This ensures that the compensatory screen is evaluated on the same decoy-normalized *z*-scale as the WT calibration.

A TCR point mutation was called compensatory for peptide mutant *p*^(*k→x*)^ if its decoy-normalized score crossed the WT reference margin:

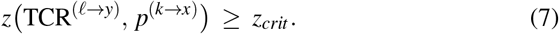

For ranking, we report both the absolute *z*-score and the improvement relative to the disrupted baseline,

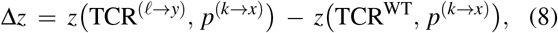

and retain the top-scoring substitutions per peptide mutant. This screen yields a concise set of single-residue hypotheses for CDR3 edits predicted to restore recognition of peptide variants that otherwise disrupt TCR binding.

### Local frustration-based validation of compensatory candidates

To assess whether modeled TCR–pMHC interfaces exhibit locally compatible energetics under single-site perturbations, we quantified residue-level interaction stability using the Frustratometer framework [21, 22]. Local energetic frustration provides a complementary metric to the RACER-m coarse-grained energy by identifying where native contacts are energetically optimized or strained within the structural context of each complex.

The Frustratometer evaluates every native residue–residue interaction by comparing its interaction energy with an ensemble of decoy interactions generated by systematically randomizing residue identities or interaction parameters while preserving the underlying geometry. The resulting frustration index classifies each contact as minimally frustrated (native energy substantially more favorable than decoys), neutral (near the decoy mean), or highly frustrated (native energy substantially less favorable).

Frustration-density profiles were computed by scanning, for each residue, the fraction of minimally frustrated, neutral, and highly frustrated interactions within a 5 Åspherical neighborhood, yielding a continuous sequence-resolved energetic map. Residue indices were partitioned into four structural regions—the MHC heavy chain (1–180), the peptide (181–190), CDR3*α* (191–389), and CDR3*β* (390–634)—and these boundaries were used when annotating frustration-density plots.

All Frustratometer analyses were performed directly on the same structural models used for RACER-m evaluation. This included the WT 1E6–ALW crystal structure (PDB 3UTT), the peptide-mutant complexes, and the predicted compensatory TCR–peptide–MHC structures. These coordinates were supplied to the Frustratometer without further refinement, ensuring that frustration metrics were computed on the exact geometries used in our modeling workflow.

## Results

We combined a structure–informed scoring model with exhaustive single–amino-acid mutagenesis to (i) quantify how peptide substitutions alter recognition by the diabetogenic 1E6 TCR (HLA-A*02:01–restricted), (ii) classify variants whose activity is retained or lost relative to the wild type (WT), and (iii) search the CDR3 sequence space for minimal TCR edits that could restore recognition on the same decoy-normalized scale.

### Structure-guided modeling recapitulates key features of the 1E6–ALW mutational scan

As a reference benchmark, we used the comprehensive single–amino-acid substitution scan of the preproinsulin epitope ALWGPDPAAA (PPI 15–24) reported for TCR clone 1E6 [18]. In this dataset, all 19 substitutions were introduced at each of the 10 peptide positions, and T-cell activation was quantified as TNF*α* release relative to the WT peptide. The scan revealed strong positional asymmetry: substitutions at the flanking residues (Ala^1^, Leu^2^, Tyr^3^, Ala^8–10^) were often tolerated, whereas mutations at the central Gly–Pro–Asp-Pro core consistently abolished activation (Figure2a). In particular, Pro^5^ and Asp^6^ residues form the dominant TCR-facing hot spot in the 1E6 structure. Although Pro^7^ strongly affects peptide–MHC stability due to its role in backbone rigidity, it does not participate in the principal TCR-facing hot spot interface [18].

We initially applied our structural biophysical framework (RACER-m [17]) to determine the extent to which ALW point mutations could be reliably predicted. Toward that end, each ALW variant was threaded onto the 3UTT complex and scored using RACER-m trained on 69 ATLAS crystal structures without exposure to the ALW mutational data. Experimental TNF responses (Figure 2a) and RACER-m predictions (Figure 2b, Supplementary Figure S1a-b) share some qualitative features in common, particularly in that the Gly^4^–Pro^5^–Asp^6^ region exhibits the largest predicted disruption, whereas most flanking substitutions lie near the WT baseline. Despite these similarities, the model had difficulty in achieving quantitative agreement with the experimental data. The overall correspondence between experiment and prediction is quantified in Figure 2c, which shows the global correlation between RACER-m *z*-scores from panel (b) and experimental TNF values from panel (a). We will return shortly to improved versions of RACER-m that greatly enhance this agreement.

**Figure 1.**
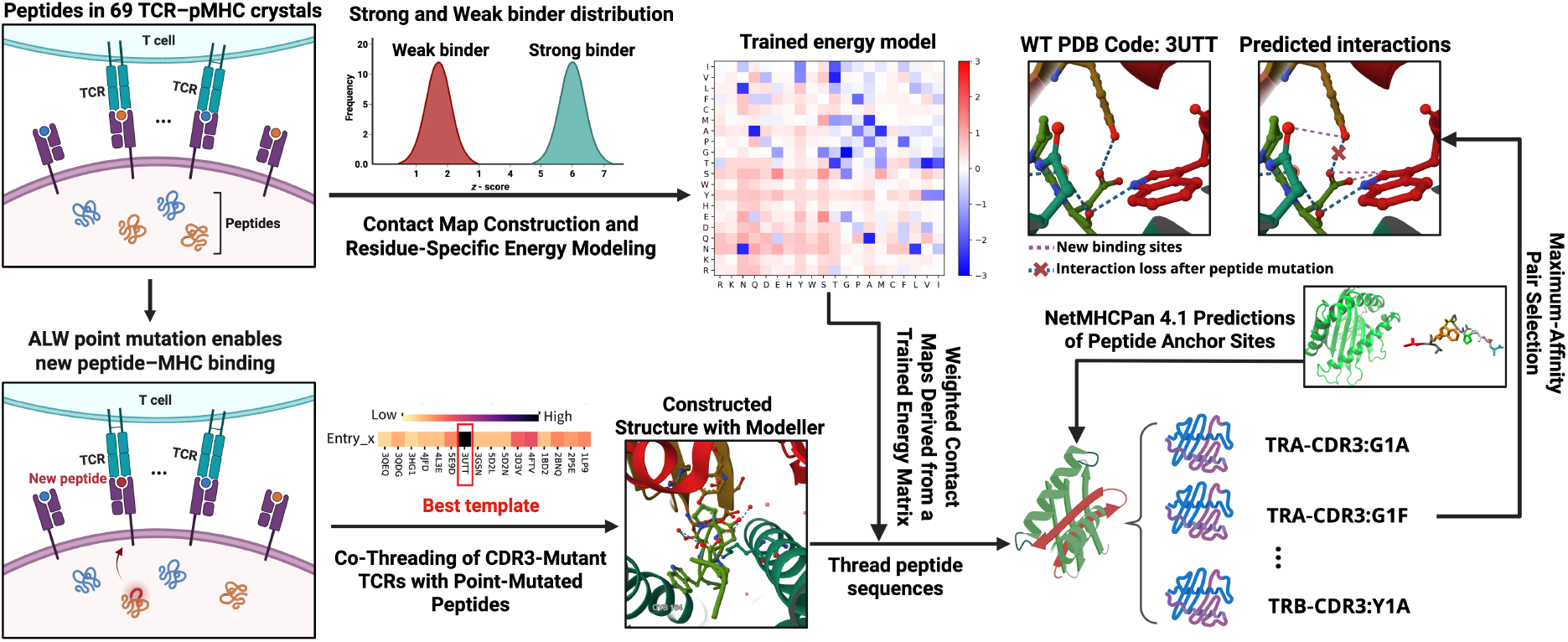
Workflow, training data, and rescue screen for 1E6–ALW using energy-based model. Training data: a curated set of 69 TCR–pMHC-I crystal structures is used to train the residue–residue energy matrix (*γ* in Eq. 1) and to define a decoy-normalized *z*-score scale for binding. **Scoring peptide mutants:** single–amino-acid substitutions are introduced into the ALW peptide and evaluated on distance-weighted contact maps with the trained model, yielding *z*-score distributions that distinguish weak vs. strong binders (top center). **Template-based modeling:** for each peptide mutant, a structural template is chosen by a composite sequence-similarity score (peptide, CDR3*α*, CDR3*β*), the complex is threaded and built in MODELLER, and energies are computed on the same decoy-normalized scale (bottom center). Example views compare the WT HLA-A*02-restricted 1E6–ALW structure (PDB: 3UTT) with a designed complex, highlighting interaction loss and potential new contacts after peptide mutation (top right). **Compensatory screen:** exhaustive single-site substitutions across 1E6 CDR3*α*/CDR3*β* are co-threaded with ALW peptide mutants classified as disrupted and subsequently re-scored; pairs that exceed the WT reference margin 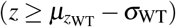 are retained and ranked as rescue candidates (right).

**Figure 2.**
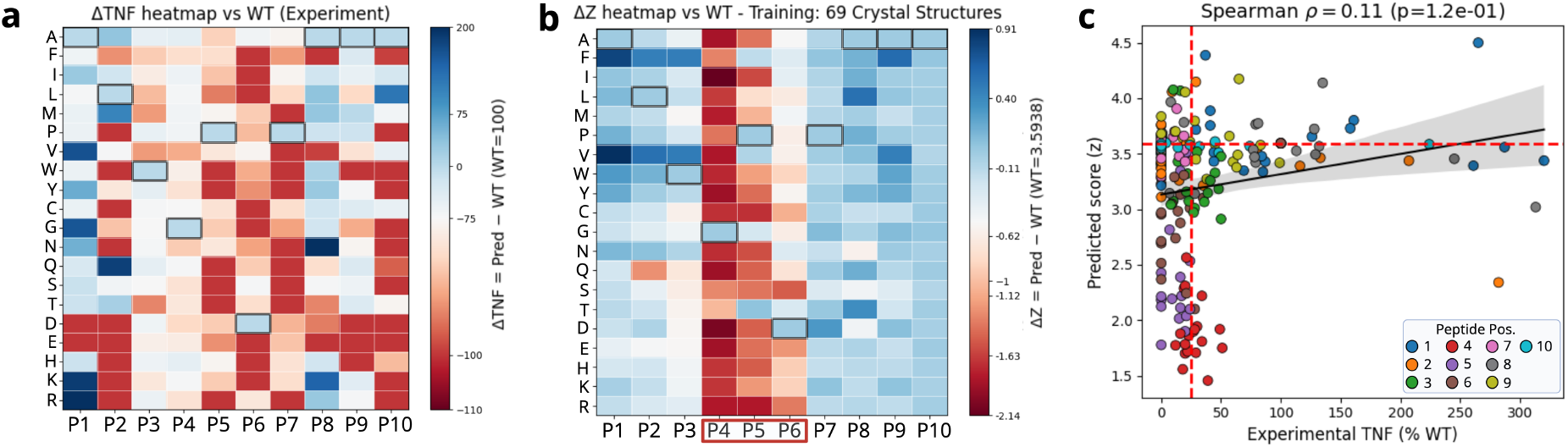
Comparison of experimental, structural, and model-predicted 1E6 responses to ALW peptide substitutions. **(a)** Experimental TNF release for all 190 single–amino–acid variants. **(b)** RACER-m predictions using the baseline 69-structure training set, capturing the central hot spot (positions outlined in red) but showing weaker agreement at flanking and MHC-facing positions. **(c)** Correlation between experimental TNF responses in panel (a) and RACER-m Δ*z* predictions in panel (b). Each point represents a single peptide substitution, colored by peptide position. The red dashed lines denote the experimental weak-activity threshold (vertical) and the WT *z*-baseline (horizontal). The fitted regression line with 95% confidence band illustrates the modest global monotonic trend between experimental activity and predicted binding score.

The modest global correlation obtained with the generic RACER-m model (Figure 2c) provides a baseline for evaluating the antigen-restricted and structurally augmented models introduced below (Figure 3e and Figure 4i). Although model specialization improves agreement with experiment, correlations between predicted scores and continuous experimental readouts remain modest. We therefore interpret RACER-m and RACER-MAP primarily in terms of their ability to recover position-resolved mutational patterns, such as the Gly^4^–Pro^5^– Asp^6^ TCR-binding hot spot, and to discriminate strong-from weak-binding variants according to Eq. (5), rather than as predictors of the precise magnitude of the experimental readout. Consistent with this interpretation, classification performance is evaluated using AUC (Figure 4j). Thus, the principal utility of the framework lies in identifying and prioritizing peptide variants according to their predicted effects on TCR recognition, rather than quantitatively reproducing experimental activation or binding levels.

**Figure 3.**
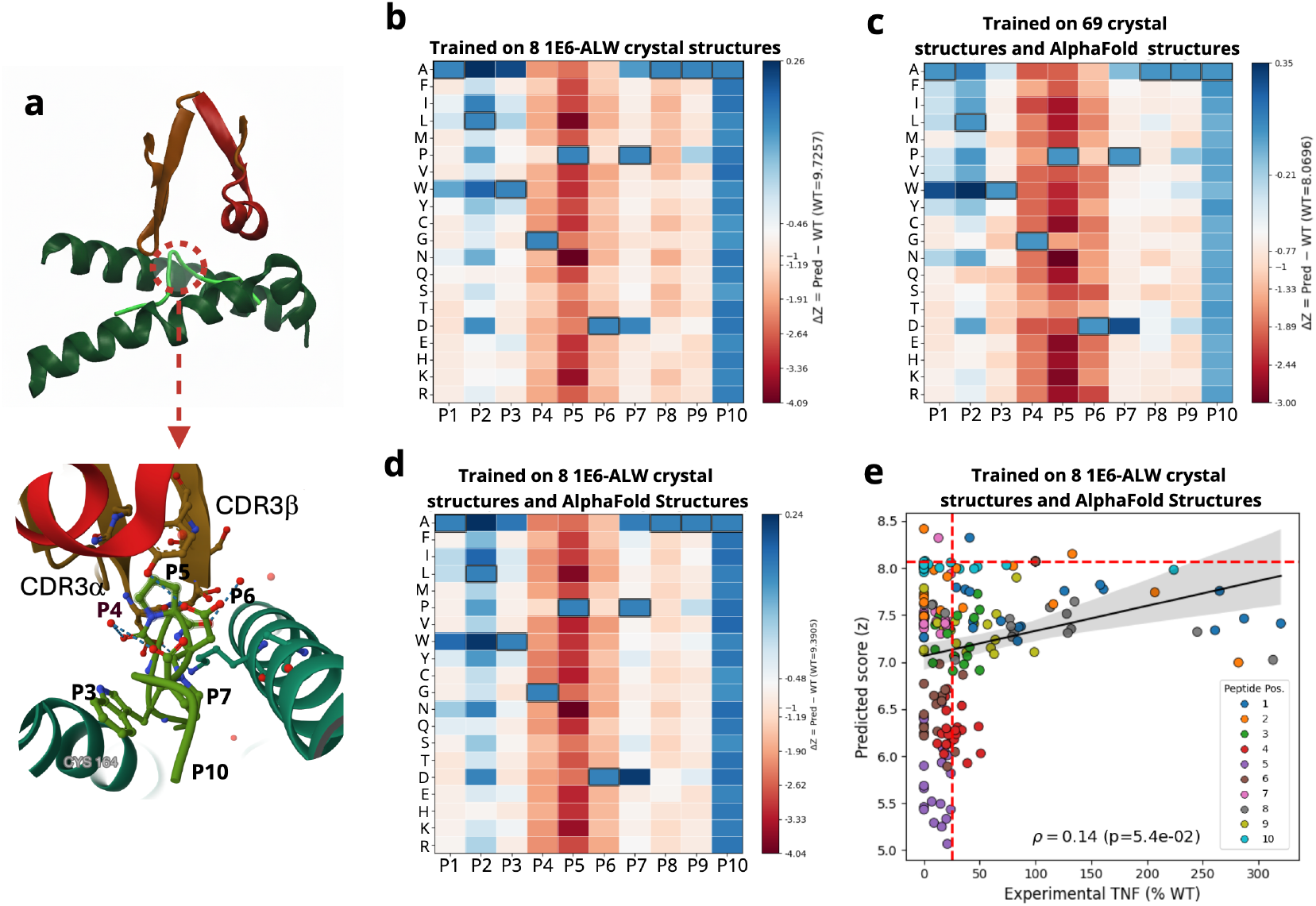
Enhanced training-set composition improves RACER-m predictions for the 1E6–ALW single–mutation landscape. **(a)** Crystallographic reference structure of the 1E6 TCR–pMHC complex (PDB: 3UTT), with the ALW peptide shown in the HLA-A*02:01 binding groove and the central Gly^4^, Pro^5^, and Asp^6^ residues highlighted to delineate the dominant TCR-facing hot-spot region. **(b)** Predicted disruption heatmap (Δ*z* = *z*_mut_*− z*_WT_) when RACER-m is trained exclusively on the eight 1E6–ALW crystal structures (*ALW*). **(c)** Predictions from the model trained on the 69 ATLAS TCR–pMHC crystal structures augmented with AlphaFold3-generated strong-binding peptide–mutant complexes (*ATLAS–AF*). **(d)** Predictions from the combined antigen-specific and structure-augmented configuration, in which RACER-m is trained on the eight 1E6–ALW crystal structures together with AlphaFold3-derived peptide–mutant complexes (*ALW–AF*). **(e)** Global correlation between experimental TNF responses and RACER-m predictions computed using the *ALW–AF* model, summarizing the quantitative agreement achieved after antigen restriction and peptide-side structural augmentation. Across all training setups, the characteristic Gly^4^–Pro^5^–Asp^6^ hot spot is consistently recovered, while incorporation of AlphaFold-derived peptide variants enhances agreement at several flanking positions.

**Figure 4.**
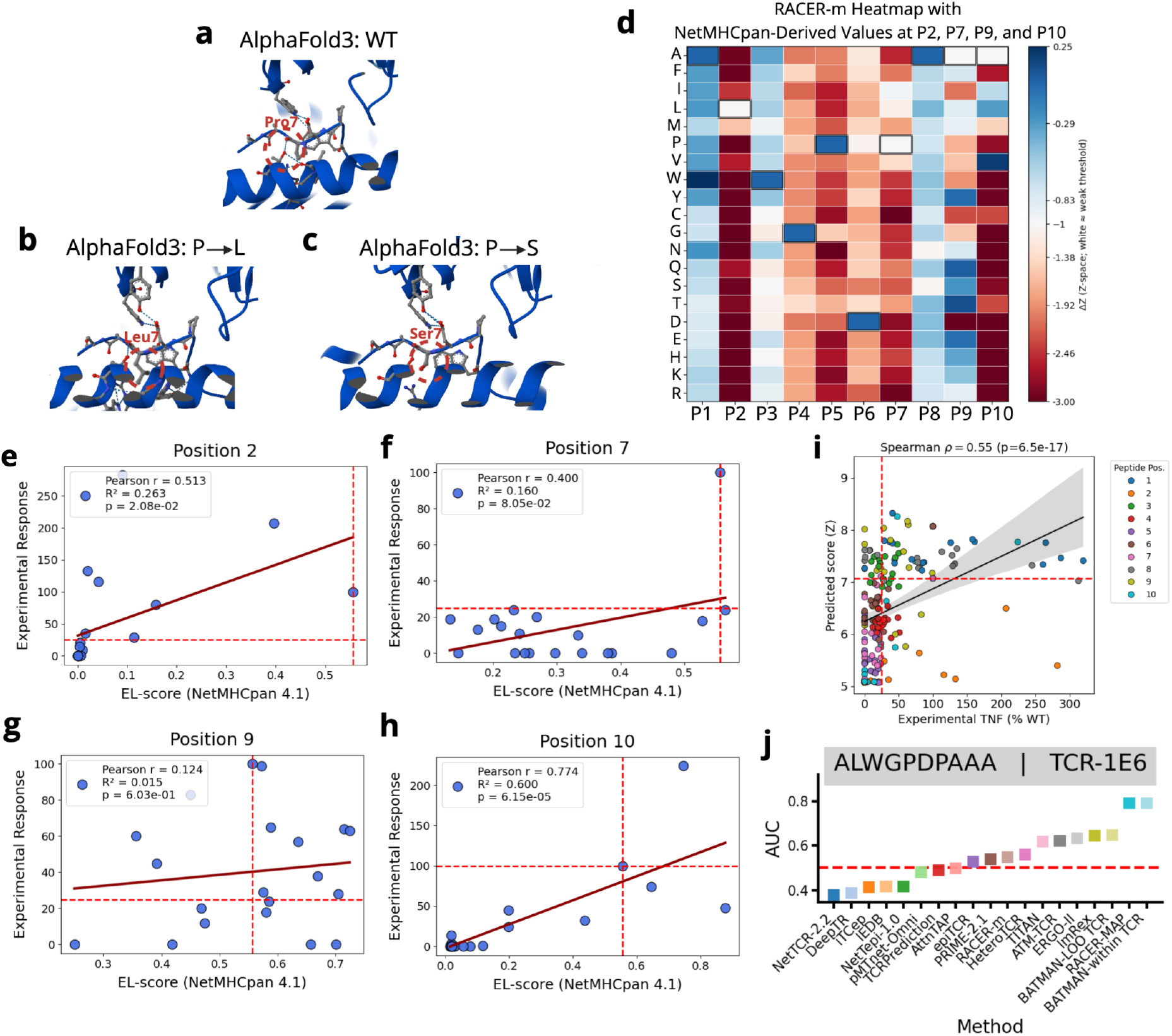
Disentangling TCR-facing and MHC-facing effects in the 1E6–ALW mutational scan. (a) AlphaFold3 model of the WT ALW peptide bound to HLA-A*02:01, showing MHC-facing versus TCR-facing orientations. (b–c) Representative Pro^7^ substitutions (Pro^7^ *→*Leu in panel b; Pro^7^ *→*Ser in panel c) illustrating how mutations reshape peptide accommodation within the MHC groove while leaving the TCR-facing surface essentially unchanged. (d) RACER-m predictions after ALW-restricted training enriched with 18 AlphaFold3-derived peptide–mutant structures; mutational effects at MHC-facing residues (Leu^2^, Pro^7^, Ala^9^, Ala^10^) are interpreted using NetMHCpan 4.1, as these positions predominantly modulate peptide–MHC stability rather than TCR contact. (e–h) Position-specific correlations between experimental TNF responses and NetMHCpan 4.1 predictions for Leu^2^ (e), Pro^7^ (f), Ala^9^ (g), and Ala^10^ (h), confirming that experimental sensitivity at these sites arises primarily from altered peptide–MHC binding. (i) Integrated correlation combining RACER-m predictions for TCR-facing positions with NetMHCpan 4.1 predictions for MHC-facing positions, demonstrating substantially improved overall agreement once peptide–MHC energetics are incorporated. (j) Benchmarking RACER-m and RACER-MAP against published TCR–pMHC specificity predictors. Classification performance (AUC) for distinguishing strong vs. weak 1E6–ALW binders among single-point peptide mutants is compared across RACER-m, RACER-MAP, and published TCR–pMHC prediction methods, including NetTCR-2.2, DeepTCR, iTCep, IEDB, NetTepi-1.0, pMTnet-Omni, TCRPrediction, AttnTAP, PRIME-2.1, HeteroTCR, TITAN, ATM-TCR, ERGO-II, ImRex, BATMAN-LOO TCR, and BATMAN-within TCR. RACER-m and RACER-MAP were trained using only 10% of the available 1E6–ALW mutational data, with the remaining 90% held out for testing. Most published comparator methods were evaluated using their existing pretrained models and were not retrained on the 1E6–ALW mutational dataset. BATMAN-within TCR was trained and evaluated using five-fold cross-validation, corresponding to approximately 80% of the 1E6-specific data for training in each fold.

### Incorporation of gain-of-function “super agonist” substitutions

Although most single–amino–acid substitutions reduce 1E6 activity, the experimental scan also contains a subset of enhancing mutations that elicit TNF responses above the WT peptide. To assess whether the specialized RACER-m model captures these gain-of-function cases, we examined the four peptide positions that contain experimentally validated super-agonist substitutions—Ala^1^, Trp^3^, and Ala^8^. For each position, we asked whether any mutant received a *z*-score exceeding the WT baseline, thereby entering the model’s strong-binder regime (Eq. 5).

Across all four positions, RACER-m correctly identified at least one experimentally enhanced mutant. For example, the A^1^ *→*V substitution, which produces the strongest enhancement experimentally, is also the top-ranked variant *in silico*. At Trp^3^, the model’s highest-scoring mutant likewise corresponds to an experimentally confirmed enhancer. Similar agreement is observed at Ala^8^, where the model consistently elevates at least one enhancing substitution above the WT.

### Biophysical model restriction to 1E6-ALW

#### Antigen-restricted training

Let us now return to addressing the issues with our initial formulation. The original RACER-m model contained only 8 1E6–ALW structures out of 69 total, and 20 strong 1E6–ALW primary pairs for binding. Because of this, we reasoned that the original learned energy matrix may not sufficiently emphasize the key contacts determining TCR specificity in ALW-related systems. To evaluate this possibility, we omitted the original TCR–pMHC structures and strong binding examples unrelated to 1E6–ALW from training. In this reduced training case, referred to as *ALW– ATLAS8*, the RACER-m model was re-trained exclusively on the eight available ALW crystal structures (WT peptide with 1E6 mutants).

Figure 3a shows the crystallographic 1E6 TCR–pMHC complex (PDB: 3UTT), with the ALW peptide displayed in the binding groove and the central Gly^4^, Pro^5^, and Asp^6^ residues highlighted to delineate the dominant TCR-facing hot-spot region. This TCR-facing hot spot constitutes the primary location of peptide discrimination by the 1E6 TCR and therefore represents the dominant source of signal utilized by the RACER-m energy model. Accordingly, the improved predictive agreement observed in Figure 3 arises not from global re-optimization of the full TCR–pMHC complex, but from more accurately emphasizing the local energetic landscape at the TCR-facing peptide interface. Structural and sequence augmentation enrich the training data specifically in the region where the TCR directly interrogates peptide side chains, sharpening the model’s sensitivity to substitutions at TCR-contacting residues while leaving MHC-facing and framework regions largely unchanged. Consequently, the observed performance gains reflect improved resolution of peptide-CDR3 interactions rather than global structural changes, consistent with the conserved overall docking geometry of the 1E6–pMHC complex.

This antigen-specific training improved agreement at both hot spot and flanking positions, demonstrating the value of context-matched templates (Figure 3b, Supplementary Figure S1c-d). As expected, the optimized energy matrix shifted from the diffuse, heterogeneous pattern learned from the full 69-structure ATLAS set to a more focused and sparsified profile (Supplementary Figure S2): strong interaction weights concentrated on a smaller set of residue pairs repeatedly observed in the 1E6–ALW interface, while irrelevant interaction modes present only in unrelated TCR–pMHC systems were suppressed. Nevertheless, substitutions at MHC-facing positions (Leu^2^, Pro^7^, Ala^9^, Ala^10^) still displayed weak correspondence with experiment.

We reasoned that the relatively sparse training set of eight available strong binding pairs was a limiting feature in training an optimized energy matrix that adequately informed binding at every position. Moreover, because none of these experimental structures include peptide substitutions, this version of the model lacked direct geometric information about how ALW mutations reshape peptide conformation. To make further progress, we next explored the role of additional sequence and structural augmentation to our training set.

#### Sequence-only augmentation with strong-binding mutants

To isolate the benefit of including additional strong binding TCR–peptide sequences without altering the structural templates, we next incorporated experimentally verified strong-binding ALW mutants at the sequence level only by randomly selecting approximately 10% of the experimentally verified strong-binding ALW mutants (TNF > 100% of WT) (Supplementary Figure S3a-b). This configuration, denoted *ALW– SB18*, improved performance modestly at tolerated positions, indicating that exposure to strong-binding sequences helps refine the learned energetic preferences. Nevertheless, because the geometry of the peptide–MHC–TCR interface remained unchanged, improvements at structurally sensitive positions were limited.

#### Structural augmentation with AlphaFold3

We therefore introduced explicit peptide-side structural diversity through the generation of AlphaFold3 models for 18 experimentally validated strong-binding mutants (TNF *>* 100% of WT), forming the *AF18* set. These *in silico* structures were incorporated into two major configurations: In the first, *AF18* was added to the full 69-structure ATLAS training set (*ATLAS–AF18*; Figure 3c), extending the baseline general model with templates diverse in TCR-ALW-mutant interactions. In the second, *AF18* was added to the antigen-restricted *ALW–ATLAS8* configuration (*ALW–AF18*; Figure 3d, Supplementary Figure S3c-d), producing an augmented training set specialized to 1E6-ALW-related point-mutant systems.

To generate the AlphaFold3-derived 1E6–pMHC complexes, we constructed mutant-specific structures by substituting the peptide sequence and the corresponding CDR3*α* and CDR3*β* loops while retaining the V*α* and V*β* framework-region sequences from the crystallographic 1E6 structure (PDB: 3UTT) and keeping the HLA-A*02:01 heavy chain and *β*_2_-microglobulin fixed. Importantly, while the V-region sequences outside CDR3 were held constant, AlphaFold3 was allowed to predict the full three-dimensional structure of each complex de novo, such that the resulting models are not constrained to reproduce the experimental 3UTT coordinates. This strategy ensured that all models differed only at the level of peptide identity and CDR3 loop sequence, while preserving a consistent receptor–MHC sequence framework across the entire panel.

Each assembled complex was submitted to the AlphaFold3 pipeline using default settings. Predicted structures were filtered by per-residue pLDDT confidence scores (*≥*85) and were further examined for backbone continuity, absence of chain detachment, and physically reasonable packing at the TCR–peptide interface. Across all 18 mutants, AlphaFold3 consistently reproduced the characteristic 1E6 docking geometry without introducing large-scale distortions, suggesting that the mutated peptides are structurally compatible with the canonical binding mode. The resulting collection of 1E6–ALW-mutant structural models formed the basis of the *AF18* augmented training configurations summarized in Figure 3.

The AlphaFold3-generated complexes should be interpreted as structural plausibility checks rather than independent validation of the predicted TCR–pMHC interactions. The 1E6– ALW–HLA-A*02:01 complex is a well-characterized crystallographic system (PDB: 3UTT), and AlphaFold3 may therefore encode structural information or closely related priors for this interaction. Consequently, the recovery of plausible docking geometries for single-point peptide variants on the same structural scaffold does not by itself establish generalization to unseen TCR–pMHC interactions. Throughout this study, we therefore use AlphaFold3-derived complexes to assess whether the predicted mutant configurations are structurally self-consistent and compatible with the RACER-m models, rather than as evidence that the corresponding variants support productive TCR–pMHC binding or T-cell activation. Such functional validation ultimately requires experimental testing.

Application of the *AF18* structures improved model performance across several non–hot-spot positions where peptide geometry plays a substantial role while also maintaining sensitivity at the central Pro^5^/Asp^6^ core. Importantly, structural augmentation provided only modest additional improvement beyond the sequence-only *ALW–SB18* model, consistent with the relatively conserved geometry of the 1E6–pMHC interface across both crystallographic and AlphaFold3-derived templates, as supported by mutual *Q* analysis comparing experimental and AlphaFold3-derived structures (Supplementary Figures S3–S5). This limited but consistent gain is further reflected at the model-parameter level: comparison of the learned energy matrices with and without AlphaFold3 augmentation (Supplementary Figure S5c) shows that structural enrichment induces localized reweighting of peptide-facing interaction terms, while leaving the overall structure of the energy matrix largely unchanged.

To quantify model–experiment agreement beyond perposition heatmaps, we compared RACER-m *z*-scores with experimental TNF values across all 190 single mutants using the *ALW–AF18* configuration. The resulting global correlation (Figure 3e) is modest — reflecting the mixture of TCR-facing and MHC-facing positions — but correctly places disruptive peptide variants below the WT z-score baseline while clustering tolerated or enhancing variants near or above WT. Position-resolved correlations in Supplementary Figure S4 reveal clearer monotonic trends, especially at the central Gly^4^–Pro^5^–Asp^6^ hot spot. Together with the per-position heatmaps in Figure 3b–d, these quantitative analyses support the improved agreement achieved after antigen restriction and peptide-side diversification.

In summary, antigen-restricted training enhances recognition of the ALW hot spot, strong-binding sequences provide meaningful improvements even without structural augmentation, and AlphaFold3-derived peptide mutants yield modest but consistent gains across structurally sensitive positions. We next turn to the discussion of mutations which primarily affect peptide–MHC binding.

### Accounting for peptide–MHC binding

Although model specialization and peptide-diverse augmentation substantially improved performance at TCR-facing positions, discrepancies remained at residues whose side chains are oriented toward the MHC rather than the TCR. This pattern reflects the structural biology of the ALW epitope: Leu^2^ and Ala^9^ constitute the primary HLA-A*02:01 anchor residues, while Pro^7^ and Ala^10^ project into the MHC floor and make minimal direct contact with the 1E6 TCR. Because RACER-m explicitly models only TCR–peptide interactions and does not include peptide–MHC energetics, substitutions at these positions are intrinsically difficult for the model to score accurately.

From a functional standpoint, the experimental readout measures T cell TNF production and therefore does not distinguish between two underlying causes of loss of activity: (i) impaired TCR recognition due to disrupted TCR–peptide geometry, or (ii) loss of peptide–MHC stability that prevents proper complex formation. Thus, weak 1E6 responses at MHC-facing positions may arise primarily from changes in MHC anchoring rather than alterations to TCR engagement.

To visualize this MHC-dominant behavior, Figure 4a–c shows AlphaFold3 structural models of WT ALW (panel a) and two representative substitutions at Pro^7^ (Pro^7^ *→*Leu, panel b; Pro^7^*→* Ser, panel c). These mutations reshape how the peptide sits within the HLA-A*02:01 groove—altering backbone geometry, pocket fit, and local peptide curvature—while leaving the TCR-facing surface essentially unchanged. Accordingly, substantial experimental changes in TNF can arise from perturbed MHC binding even when the TCR-contacting residues retain WT-like configuration, explaining why RACER-m systematically underperforms at these sites.

To interpret these MHC-driven effects, we incorporated NetMHCpan 4.1 predictions [20] for peptide–MHC binding affinity and overlaid them with RACER-m predictions obtained after ALW-restricted training enriched with 18 AlphaFold3-derived peptide–mutant structures. In this configuration (Figure 4d), substitutions at the four MHC-facing positions (Leu^2^, Pro^7^, Ala^9^, Ala^10^) are interpreted using NetMHC-pan 4.1 because their experimental effects arise primarily from altered peptide–MHC stability, whereas all TCR-facing positions retain RACER-m predictions. As expected, these MHC-facing sites show the largest NetMHCpan 4.1-predicted shifts in binding affinity, in sharp contrast to the central Gly^4^– Asp^6^ hot spot, where mutations minimally affect MHC engagement. Experimental TNF responses correlate strongly with NetMHCpan 4.1 predictions across all four MHC-facing positions (Figure 4e–h), confirming that the observed experimental sensitivity at these residues is driven predominantly by peptide–MHC stability rather than changes in TCR recognition.

Together, these analyses establish a mechanistic framework for interpreting the mixed sources of signal in the ALW mutational scan: RACER-m accurately captures TCR-facing interactions, whereas NetMHCpan 4.1 accounts for positions dominated by allele-specific peptide anchoring. By combining these complementary components, the integrated correlation shown in Figure 4i—the composite model we hereafter refer to as RACER-MAP (Methods)—demonstrates that model–experiment agreement improves substantially once MHC-driven effects at Leu^2^, Pro^7^, Ala^9^, and Ala^10^ are interpreted through peptide–MHC stability rather than TCR–peptide contacts. This unified view provides a complete explanation for the positional asymmetry observed experimentally and motivates our subsequent use of the TCR-focused model to identify compensatory receptor edits capable of restoring recognition of escape variants.

We additionally benchmarked classification performance (AUC for distinguishing strong vs. weak 1E6–ALW binders) against a cohort of published sequence- and structure-based TCR–pMHC specificity predictors, situating RACER-m and RACER-MAP within the broader specificity-prediction literature (Figure 4j). RACER-m/RACER-MAP were trained on only 10% of the 1E6–ALW mutational data, whereas the published comparator methods were trained on 80% of the data, with the exception of BATMAN-LOO TCR, which used a leave-one-out scheme—trained on all other TCRs in the BATMAN reference cohort and tested on 1E6 as the single held-out clone—rather than a within-TCR data split at all [8]. Despite this pronounced data disadvantage, baseline RACER-m achieved an AUC of 0.5491, while RACER-MAP (AUC = 0.7914) and BATMAN-within TCR (AUC = 0.7915) ranked among the top-performing methods in the cohort, and sequence-only deep-learning classifiers (e.g., NetTCR-2.2, DeepTCR, ImRex) spanned a wide range of intermediate performance (AUC*≈* 0.3772–0.6480). These results indicate that, while the coarse-grained RACER-m energetics alone are only weakly discriminative for this system, augmenting with structural contact information (RACER-MAP) meaningfully closes the gap with specialized published classifiers trained on substantially more 1E6-specific data.

Together, Figures 3 and 4 establish a clear separation of roles: RACER-m accurately captures the TCR-facing component of peptide recognition, while NetMHCpan 4.1 accounts for experimental sensitivity driven by peptide–MHC anchoring, and their combination (RACER-MAP) yields the integrated, structure- and MHC-aware prediction used throughout the remainder of this study. This decomposition resolves the apparent positional asymmetry in the ALW mutational scan and, critically, isolates the portion of the binding signal that is explainable by TCR-peptide interactions. Building on this separation, we next leverage the TCR-focused model to systematically predict compensatory edits to the 1E6 receptor capable of restoring recognition of disruptive antigenic variants.

### Compensatory TCR mutations predicted to restore recognition of escape antigenic variants

Having established a unified explanation for the positional asymmetry in the ALW scan — where TCR-facing effects are captured by RACER-m and MHC-facing effects by NetMHC-pan 4.1 — we next asked whether the TCR itself can compensate for disruptive peptide mutations. Specifically, we examined whether single–site edits to the 1E6 CDR3 loops could restore recognition of experimentally weak ALW variants. We focused on escape variants that were categorized as weak both empirically (TNF *≤*25% of WT) and in our predictions (Eq. 5), yielding 115 SAPVs across the ten peptide positions. For each escape peptide, we exhaustively enumerated single amino–acid substitutions in CDR3*α* and CDR3*β* and re-scored the cognate pMHC with RACER-m. A TCR edit was deemed <u>compensatory</u> if the resulting complex fell within the strong–binder distribution, i.e., *z≥ z*_crit_ as defined in Eq. 5.

The structural context for compensatory mutagenesis is summarized in Figure 5. Figure 5a shows the WT CDR3*α* contact map, with the central Gly^4^–Pro^5^–Asp^6^ hot spot highlighted in black and the surrounding flanking positions annotated in purple. Figure 5b presents the corresponding WT CDR3*β* contact map using the same color scheme. Together, these maps illustrate that CDR3*α* forms the dominant set of peptide-facing contacts at the hot spot, while CDR3*β* provides additional stabilizing interactions that support cooperative recognition of the ALW peptide.

**Figure 5.**
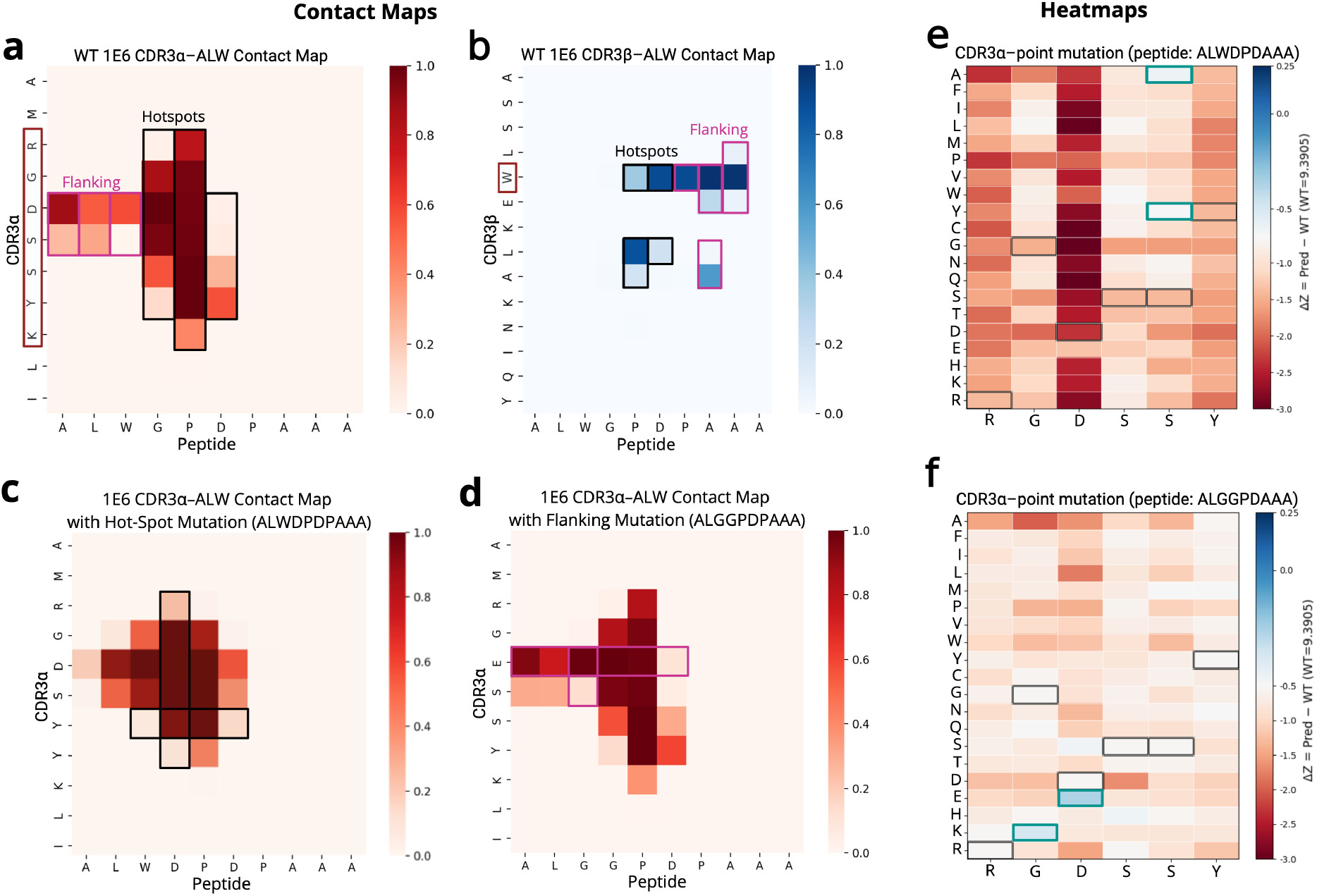
Predicted compensatory TCR mutations restoring recognition of escape variants. (a) WT CDR3*α* contact map for the 1E6–ALWGPDPAAA TCR–pMHC complex (PDB: 3UTT), with the peptide hot spot highlighted in black and the flanking region in purple. (b) WT CDR3*β* contact map with the same annotations. (c) Compensatory rescue at a hot-spot site: the Gly^4^ *→*Asp escape variant is predicted to be restored by the CDR3*α* substitution S^96^ *→*Y, which introduces new favorable peptide–TCR contacts. (d) Compensatory rescue at a flanking site: for the peptide variant `ALGGPDPAAA`, the CDR3*α* substitution D *→*E restores the disrupted interaction pattern and recovers recognition. (e) Heatmap of compensatory potential across CDR3*α* substitutions for the case in panel (d). (f) Heatmap of compensatory potential across CDR3*α* substitutions for the case in panel (e). In panels (e,f), blue blocks denote successful rescues (strong-binder predictions), and green squares highlight the specific positions where compensatory substitutions were identified.

A representative hot-spot case is the Gly^4^ *→*Asp escape mutant, which experimentally abolishes 1E6 activation. This substitution disrupts the WT interface and yields a weak-binding complex. RACER-m predicts that an *α*-chain S^96^ *→*Y edit restores recognition by forming new favorable contacts with the mutated Asp side chain, shifting the complex back into the strong-binder regime (Figure 5c). This illustrates a hot-spot escape variant that can be compensated through a single CDR3*α* substitution.

Compensation is also observed at flanking sites. For example, the peptide variant `ALGGPDPAAA` reduces recognition experimentally, yet RACER-m predicts that a D *→*E substitution in CDR3*α* restores favorable contacts and recovers binding (Figure 5d). These cases illustrate that peptide flanks are structurally more tolerant, making them high-probability candidates for compensation through single-site TCR edits.

We next examined escape mutations at positions where both CDR3*α* and CDR3*β* contribute to peptide recognition. A representative example is a substitution at Pro^5^, one of the central hot-spot residues. In this case, RACER-m predicts that successful rescue requires cooperative adjustments from both chains, as shown in Figure S6 and Figure S7 and further detailed in Supplementary Section *Dual-contact compensatory regime at the Pro*^5^ *peptide hot-spot*. This dual-contact regime is less amenable to single-site edits, reflecting the indispensable structural role of Pro^5^ in stabilizing the TCR–peptide interface.

To systematically evaluate compensatory potential, we generated heatmaps of rescue probability across all amino–acid substitutions (Figures 5e,f). Panel 5e corresponds to the Gly^4^ *→*Asp escape case (previously shown in panel 5c). Here, two CDR3*α* substitutions—S*→* A and S*→* Y—were identified as compensatory, each restoring the complex to the strong–binder regime. Panel 5f corresponds to the flanking variant `ALGGPDPAAA` (shown structurally in panel 5d). In this case, RACER-m predicts a single compensatory edit, D*→* E in CDR3*α*, aligned with the experimentally observed tolerance at this position.

The dual-contact Pro^5^ case, where both CDR3*α* and CDR3*β* interact with the peptide, is analyzed separately in Supplementary Figure S6. Rescue in this regime is markedly constrained: several CDR3*α* edits (S *→*Q, K*→* Q, S*→* V) restore binding only when CDR3*β* remains unaltered, and true cooperative rescue occurs in only one combination (CDR3*β* W*→* L together with CDR3*α* S*→* Y; Supplementary Figure S7).

Together, Figures 5c–f and Supplementary Figure S6-S7 reveal three compensatory regimes: (i) flanking positions that are readily rescued by single CDR3*α* edits (panel 5f), (ii) hotspot escape variants remediated primarily by CDR3*α* substitutions (panel 5e), and (iii) dual-contact hotspot mutations that require coordinated CDR3*α*/CDR3*β* editing and exhibit limited rescue potential (Supplementary Figure S6-S7).

### Computational plausibility checks for predicted compensatory complexes

To evaluate the robustness and physical plausibility of RACER-m–predicted compensatory complexes, we performed two complementary analyses: (i) an independent structural plausibility check using AlphaFold3 [23], and (ii) an energetic plausibility check using the Frustratometer [21, 22].

#### Structural plausibility check using AlphaFold3

To assess whether energetic restoration is accompanied by geometrically plausible interfaces, we modeled the full compensatory TCR–pMHC complexes using AlphaFold3. The resulting interfacial contact maps (Figure 6a–b) closely resemble those derived from RACER-m (Figure 5c,d), supporting the structural plausibility of the predicted compensatory interactions. This agreement should be interpreted as consistency between two computational structure-generation approaches rather than as experimental confirmation of productive TCR–pMHC binding.

**Figure 6.**
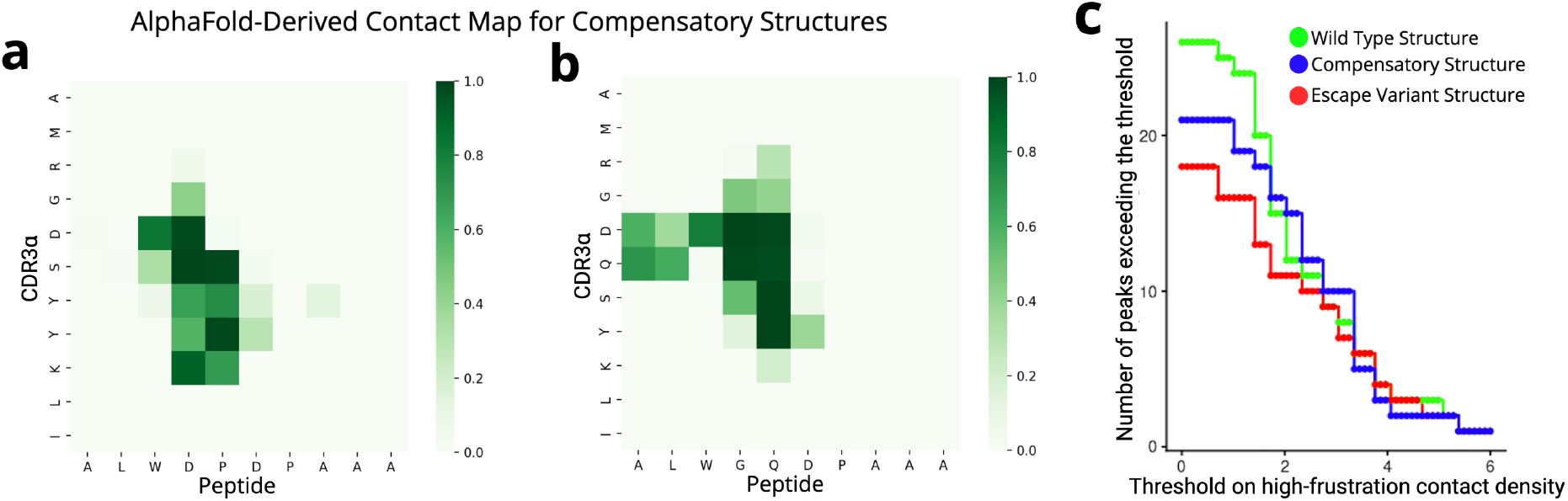
Structural and energetic validation of predicted compensatory complexes. **(a**,**b)** AlphaFold3-derived interfacial contact maps for compensatory 1E6–pMHC complexes. **(a)** Hot-spot escape mutant Gly^4^ *→*Asp rescued by the CDR3*α* substitution S^94^ *→*Y. **(b)** Flanking escape variant `ALGGPDPAAA` rescued by the CDR3*α* substitution D *→* E. In both cases, AlphaFold3 reproduces the peptide-facing contact patterns predicted by RACER-m, confirming that the compensatory solutions correspond to geometrically feasible TCR–peptide interfaces. **(c)** Number of low-frustration peaks exceeding a given threshold. The number of local maxima (“peaks”) in the minimally frustrated (low-frustration, green) contact–density profile is plotted as a function of the applied threshold for the wild-type (green), compensatory (blue), and escape (red) complexes. Each peak corresponds to a contiguous cluster of energetically favorable, minimally frustrated interactions along the TCR–peptide interface (see Figure S10-S12 for representative frustration–density profiles and peak identification). At low thresholds, the wild-type structure exhibits the largest number of low-frustration peaks, indicating a broadly distributed network of stabilizing contacts. As the threshold increases, the number of surviving peaks decreases in all cases, with the compensatory complex retaining fewer but more localized low-frustration peaks, consistent with energetic redistribution rather than full restoration of the wild-type stability pattern.

For the hot-spot escape mutant Gly^4^ *→*Asp compensated by CDR3*α* S^96^ *→*Y (Figure 6a), AlphaFold3 recapitulates the restored CDR3*α*–peptide contacts predicted by RACER-m, including new interactions with the Asp side chain. For the flanking variant `ALGGPDPAAA` compensated by CDR3*α* D*→* E (Figure 6b), the AlphaFold3 map similarly preserves the peripheral contact pattern that supports recovered binding. Additional AlphaFold3 maps for the dual-contact Pro^5^ case are provided in Supplementary Figure S9.

#### Energetic validation using the Frustratometer

The Frustratometer quantifies local energetic favorability by comparing the native interaction energy of each residue pair with energies drawn from an ensemble of randomized decoys. Contacts with substantially lower native energy are classified as minimally frustrated (green), indicating stable, well-packed interactions; contacts near the decoy average are neutral (gray); and contacts with higher-than-expected energies are highly frustrated (red), reflecting strained or suboptimal interactions. Thus, low frustration corresponds to energetically favorable and stabilizing interactions, whereas high frustration indicates reduced local stability at the interface. To characterize how stabilizing interactions are spatially organized, we computed residue-resolved contact–density profiles of minimally frustrated interactions along the TCR interface. Local maxima (“low-frustration peaks”) in these profiles correspond to contiguous clusters of minimally frustrated contacts, reflecting energetically stable regions that act as structural anchors within the interface (Figure S10-S12).

To quantify how frustration is distributed along the interface, we analyzed the number of local peaks exceeding a given threshold in the low-frustration contact–density profile as a function of an increasing threshold (Figure 6c), focusing on the CDR3*α* region. Here, the per-residue contact–density profile reports the spatial concentration of minimally frustrated (stabilizing) contacts; a peak exceeding a specified threshold therefore identifies a residue position that participates in a locally stable interaction network. At low thresholds, the WT 1E6–ALW complex exhibits the largest number of low-frustration peaks, consistent with a broadly distributed energetic landscape across the interface. As the threshold increases, weaker peaks are progressively eliminated in all cases; notably, the escape variant shows an overall reduction in surviving peaks, reflecting a collapse of stabilizing interactions. By contrast, the compensatory complex retains fewer but more pronounced high-intensity peaks at higher thresholds, indicating that compensatory mutations redistribute stabilizing interactions into a smaller number of focused energetic hotspots rather than reconstituting the full WT frustration topology. The corresponding analysis for the CDR3*β* region is shown in Supplementary Figure S10a–b, where the peak– threshold curves largely overlap across all variants, consistent with the more conserved energetic role of this region.

Consistent with this global behavior, the residue-resolved frustration–density profiles for the WT, escape variant, and compensatory complexes (Supplementary Figure S10c–e) are highly similar. The gray dashed vertical demarcation lines partition the sequence into its structural regions—MHC heavy chain (residues 1–180), ALW peptide (181–190), CDR3*α* (191–389), and CDR3*β* (390–634)—enabling direct comparison across domains. In the WT complex, most TCR– peptide contacts are minimally frustrated, indicating a well-optimized interface. Following introduction of the compensatory S94 *→* Y substitution in CDR3*α*, the predicted structure preserves this regional organization, restores the balance between minimally and highly frustrated contacts in the CDR3*α* region, and does not introduce new clusters of high frustration at the interface. Complementary contact maps and frustration– density distributions for CDR3*α* across all three cases are shown in Supplementary Figure S12, while the corresponding analyses for CDR3*β* are provided in Supplementary Figure S13.

At the residue level, mutational frustration analysis of CDR3*α* position 94 (Supplementary Figure S13) provides further insight into the energetic basis of compensation. Each column corresponds to a residue contacting position 94, and the letters within each column report the frustration index for all possible amino-acid substitutions at this site. In the context of the Gly^4^ *→*Asp escape peptide, Ser94 lies in a neutral or mildly frustrated regime, consistent with its inability to stabilize the altered interface. In contrast, tyrosine consistently occupies the minimally frustrated region, indicating that the S94 *→*Y substitution alleviates local energetic strain and forms more compatible interactions with the Asp-substituted peptide. Together, these analyses show that compensatory mutations restore a WT-like energetic landscape by concentrating stabilizing interactions at key interface positions.

Together, the Frustratometer and AlphaFold3 analyses indicate that RACER-m–predicted compensatory mutations yield interfaces that are both energetically and structurally consistent with the WT strong-binding state. Rather than creating a new binding mode, the compensatory edits restore a WT-like interaction network in the context of an escape peptide, offering a computational, plausibility-level argument for functional rescue of recognition.

We reiterate that this argument rests entirely on computational plausibility checks (AlphaFold3 structural modeling and Frustratometer energetics), both applied to structures generated in silico, and does not by itself demonstrate that any specific proposed CDR3 edit restores WT-like TCR–pMHC interactions or T-cell activation in a real biological system.

Combined with the small size of the crystallographic training set underlying RACER-m for this system (eight 1E6–ALW structures; see *Antigen-restricted training*), which raises a legitimate concern about overfitting and robustness even where AlphaFold3-derived structures are used to augment training, we consider the specific compensatory CDR3 edits proposed here as testable hypotheses for future experimental follow-up rather than validated rescue mutations.

To further assess the robustness of the underlying scoring framework beyond the 1E6–ALW system, we applied the same antigen-restricted, structurally augmented workflow to two independent TCR–pMHC systems (see Supplementary Information, *Generalization of the 1E6–ALW Workflow to Independent CMV and SARS-CoV-2 TCR–pMHC Systems*; Figure S14). For the CMV-specific TCR81-14 clone, RACER-MAP trained using only 20% of the available clone-specific mutational data achieved an AUC of 0.786, comparable to the top-performing BATMAN-within-TCR model (AUC = 0.791), which used approximately 80% of the clone-specific data for training in each cross-validation fold. For the SARS-CoV-2-specific SVAR-73 clone, RACER-MAP trained using only 10% of the available mutational data together with the 163-structure ATLAS dataset, which contains no SARS-CoV-2 TCR–pMHC examples, achieved an AUC of 0.834. Although these independent benchmarks do not experimentally validate the specific compensatory CDR3 mutations proposed above, they provide additional evidence that the antigen-restricted, structure-augmented RACER-MAP framework retains predictive performance beyond the 1E6–ALW system.

## Discussion

The reliable identification of TCR-pMHC recognition making (or breaking) single amino acid variants is a difficult problem, originating from the fact that point mutated amino acid sequences often behave very differently from their nearly identical WT analog [24, 25]. This study integrates exhaustive single–amino–acid mutagenesis with a structure–informed, coarse–grained energy model to probe how the diabetogenic 1E6 TCR recognizes its HLA-A*02:01–restricted ligand and how specific substitutions disrupt or enhance this interaction. Without residue–specific re–fitting, the model captured the dominant qualitative features of this highly peptide–centric interface, including the critical dependence on central peptide residues, broad tolerance at the N– and C–terminal flanks, and strong positional asymmetry in mutational sensitivity. These results highlight the capacity of a compact, physics-guided scoring framework to approximate the mutational landscape of a TCR–pMHC complex.

The model recapitulates prior structural findings that central peptide residues act as energetic hot spots, whereas terminal positions contribute less directly to recognition. By coupling residue–level interaction potentials with a decoy-normalized scoring scheme, RACER-m was able to distinguish disruptive from tolerated substitutions and identify gain-of-function variants. Use of this biophysical approach as a decision framework provides a practical means of mapping mutational tolerance, with direct applications in antigenic escape surveillance and neoantigen prioritization.

At the same time, the analysis revealed clear limits of the baseline training. While 1E6 TCR specificity predictions on the ALW WT and SAPV peptides aligned closely with the experimental findings at the central Gly^4^–Pro^5^–Asp^6^ cluster, agreement was poor at the MHC anchor residues (Leu^2^, Ala^9^) and at Ala^10^. These discrepancies stem from biological and methodological factors. Positions 2 and 9 serve as canonical MHC anchors that primarily stabilize peptide binding to HLA-A*02:01 rather than forming direct TCR contacts; consequently, mutations at these sites may modulate TCR activation indirectly, but such effects are not explicitly represented in RACER-m. Position 10 lies outside the canonical 9-mer binding register, meaning that its contributions are not effectively captured by the coarse-grained interface potential. Together, these features explain the lack of correspondence between experiment and model at positions 2, 9, and 10. By contrast, incorporating one representative peptide mutant structure generated with AlphaFold3 improved concordance at tolerated flanking positions while preserving sensitivity to the central hot spot, underscoring the value of targeted augmentation for peptide-side coverage. More generally, the addition of structurally diverse complexes and subsequent reoptimization reduces statistical noise arising from sparse sampling, while simultaneously introducing new or previously under-weighted contacts into the training set. This process can shift model emphasis toward residue–residue interactions that are physically meaningful but underrepresented in the original dataset, thereby allowing residue–residue interaction terms that were poorly sampled in the original training set to acquire statistically meaningful weights. In this way, targeted structural augmentation refines the interaction landscape without altering the underlying functional form of the model.

The same framework was extended to probe whether single TCR substitutions could restore recognition of experimentally defined escape variants. By exhaustively scanning CDR3*α* and CDR3*β*, RACER-m identified compensatory mutations that successfully rescued recognition for a subset of escape peptides. These edits often clustered at paratope sites facing the peptide hot spot, demonstrating that even single-site modifications in the TCR can restore functional binding in cases where peptide substitutions disrupt recognition. This observation highlights both the adaptability of the TCR–peptide interface and the utility of structural biophysical modeling for the rational identification of therapeutically relevant TCRs.

The predicted compensatory mutations should be regarded as computationally supported hypotheses rather than experimentally validated rescue mutations. AlphaFold3 and Frustratometer analyses establish structural and energetic plausibility, but do not demonstrate restoration of TCR–pMHC binding or T-cell activation. Moreover, because AlphaFold3 is used both for structural augmentation and for evaluating predicted compensatory complexes, agreement between these analyses is not fully independent.

Several limitations remain. The current framework is based on restrained, threaded complexes and a fixed residue–pair contact window, without explicit modeling of backbone dynamics, side-chain flexibility, or long-timescale conformational sampling. It also omits contextual features such as coreceptor interactions, post-translational modifications, and the role of the broader TCR repertoire. This analysis focused on the response of a single TCR against associated peptide point-variants, and the antigen-restricted, structurally augmented workflow was benchmarked based on antigen specificity prediction for independent CMV- and SARS-CoV-2-specific TCRs independent systems. Our findings support the value of this approach in the setting of limited clone-specific training data for other immunological contexts. Subsequent analysis and validation on a broader class of systems, in addition to prediction of multiple TCR systems simultaneously, provide future opportunities to extend these methods. In particular, flexible ensemble modeling, hybrid physics–machine learning scoring, and integration with rapidly expanding structural datasets could substantially enhance predictive power. Future work will extend our methods to include applications to additional experimental datasets across diverse TCR–pMHC contexts, specifically focusing on cases where both antigen and TCR are varied. These will be important for assessing how broadly this modeling strategy can be applied. This work demonstrates that a biophysics-informed model can capture determinants of TCR recognition and immune escape, thereby providing a framework for predicting T cell responses. Continued refinement and expansion to additional systems will help to advance precision immunology by linking mutational landscapes to therapeutic design and the control of immune escape.

## Data Availability

All data needed to evaluate the conclusions in the paper are present in the paper and/or the Supplementary Materials. All scripts and input files needed to reproduce the results are publicly available via Zenodo (RACER-multi_template v1.0.0; DOI: 10.5281/zenodo.8374294).

## Author contributions

ZSG, HL, XL, JNO, JTG designed the study. ZSG performed the research. ZSG, HL, XL, JNO, JTG analyzed the results. ZSG, HL, XL, JNO, JTG wrote the manuscript. JTG supervised the research. All authors approve of the final manuscrip version.

## Acknowledgment

HL, JNO, and JTG acknowledge support from the NSF Physics Frontier Center program via grants NSF-PHY2019745 and PHY-2607619. JTG was supported by the Cancer Prevention Research Institute of Texas (RR210080) and the National Institute of General Medical Sciences of the NIH (R35GM155458). JTG and JNO are CPRIT Scholars in Cancer Research.

