## Supplementary material for "A biophysical framework to accurately identify antigen single-amino acid escape variants and corresponding variant-specific compensatory TCR sequences": SI

### Per-position agreement under baseline and ALW-restricted training

To evaluate how well RACER-m predictions recapitulate the experimental mutational scan of ALWGPDPAAA [1], we first considered the baseline model [2] trained on 69 crystal structures from the ATLAS dataset. Figure S1 summarizes the position-wise comparison between predicted $z$-scores and experimental TNF responses. Panel a shows the per-position correlation values across the 10 peptide sites, while panel b reports the corresponding per-position classification agreement (retained vs. weak). As described in the main text, the model captures the central hot spot (positions 4–6), but shows weaker agreement at the peptide flanking regions, resulting in limited coverage of peptide mutational diversity in the baseline training set.

We next trained RACER-m with structural restriction to the eight previously reported crystal structures of the ALWGPDPAAA peptide in complex with 1E6 TCR variants (WT peptide, mutated TCR). Figure S1c,d shows the corresponding position-wise analysis. Panel c reports per-position correlation values, and panel d the classification agreement. Compared with the 69-structure baseline, ALWGPDPAAA-only training yields stronger position-wise concordance with experiment, indicating that antigen-specific structural templates improve prediction fidelity.

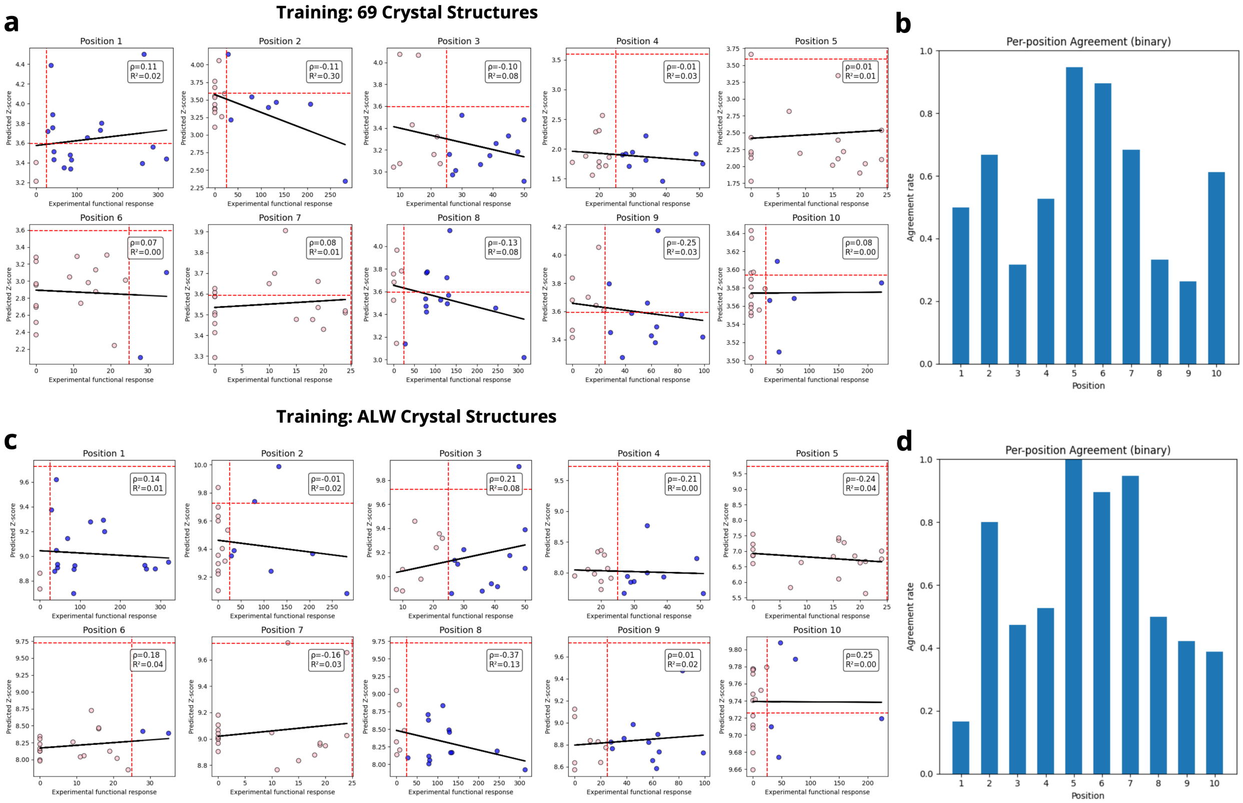


Figure S1: Per-position comparison between experimental activation and RACER-m predictions. (a) Per-position correlation with training on 69 ATLAS crystal structures. (b) Per-position classification agreement for panel a. (c) Per-position correlation with training restricted to eight ALW peptide crystal structures. (d) Per-position classification agreement for panel c.

### Learned energy matrices under alternative training regimens

The learned pairwise interaction matrices provide a concise view of how RACER-m encodes residue–residue energies under different training regimens. Figure S2a depicts the energy matrix obtained from training on the full 69-structure ATLAS dataset, while Figure S2b gives the energy matrix resulting from the antigen-restricted *ALW–ATLAS8* training set. While these matrices share some overlap, significant differences can be appreciated in large part owing to couplings concentrated on residue pairs characteristic of the 1E6-ALW interface. This distinction highlights how context-matched training enables the model to encode biological context-specific interactions.

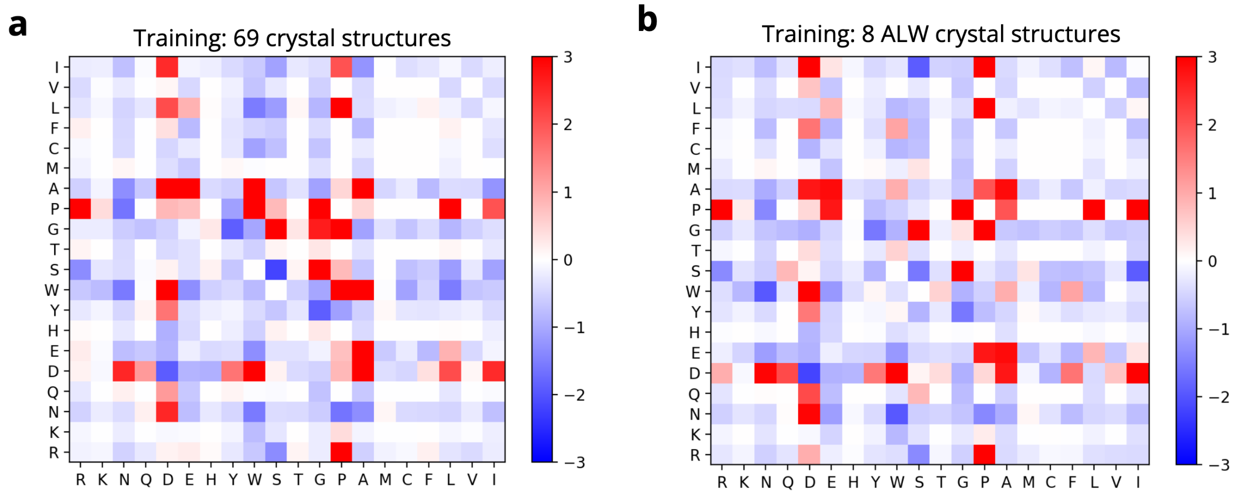


Figure S2: **Impact of training-set restriction on the RACER-m energy matrix.** **(a)** Interaction matrix learned from the full 69-structure ATLAS set shows diffuse, heterogeneous coupling patterns reflecting diverse, unrelated TCR specificities. **(b)** Matrix learned from the antigen-restricted *ALW–ATLAS8* training set becomes noticeably more focused, with strong weights concentrated on a small set of residue pairs characteristic of the 1E6–ALW interface. Restricting training to context-matched structures therefore sharpens the learned energetic features relevant for 1E6 recognition.

### Per-position agreement following strong-binder sequence enrichment

To mitigate the imbalance of peptide coverage in the training set, we randomly selected 18 experimentally validated strong-binder peptide mutants and added their sequences to the training data. These additional sequences were incorporated in two settings: (i) training on 69 ATLAS crystal structures together with 18 strong-binder peptide sequences, and (ii) training on 8 ALW crystal structures together with the same 18 strong-binder peptide sequences. Figure S3 summarizes the results. Panel a shows per-position correlation for the 69 ATLAS plus 18 strong-binder setting, and panel b the corresponding classification agreement. Panel c shows per-position correlation for the 8 ALW plus 18 strong-binder setting, and panel d the corresponding classification agreement. In both cases, enriching the training set with strong-binder sequences improved concordance at flanking sites while retaining sensitivity to the $Pro^5/Asp^6$ hot spot.

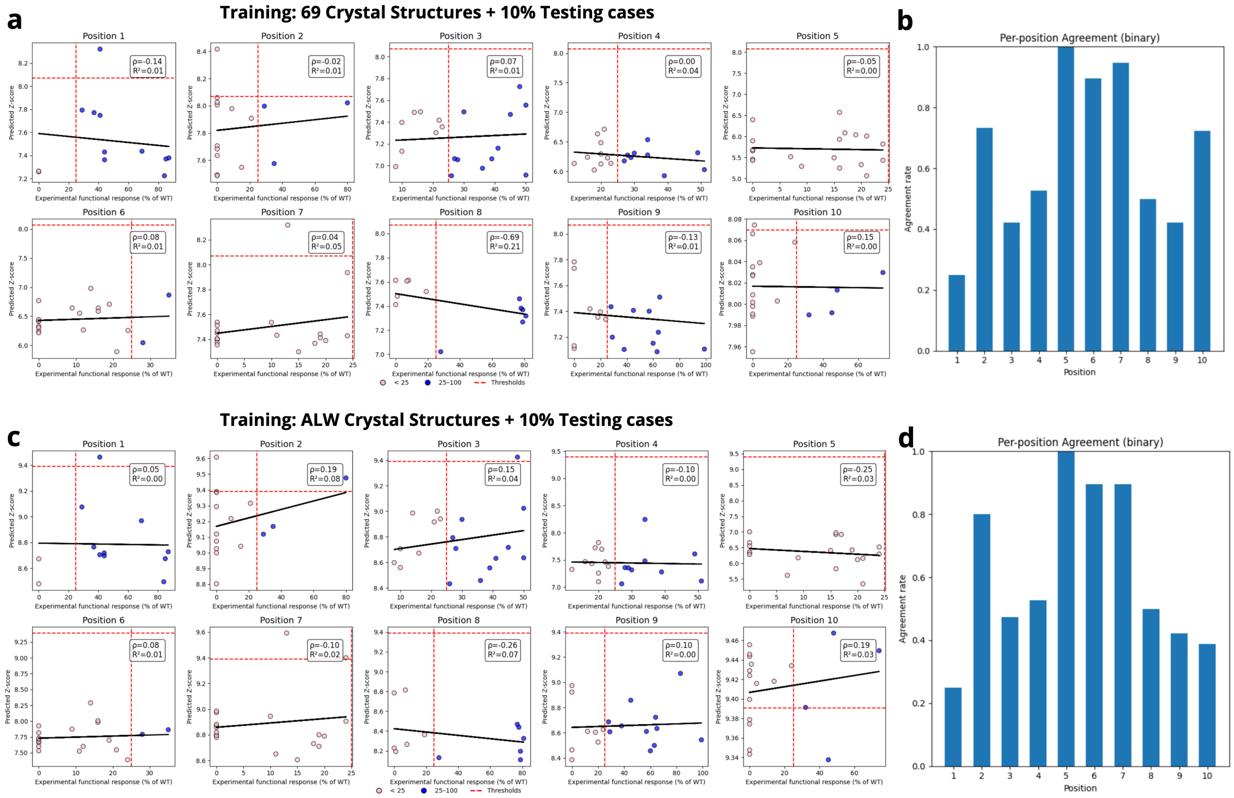


Figure S3: Per-position comparison of RACER-m predictions with experimental activation after enriching the training set with 18 experimentally validated strong-binder peptide sequences. (a) Per-position correlation with training on 69 ATLAS crystal structures together with 18 strong-binder sequences. (b) Per-position classification agreement for panel a. (c) Per-position correlation with training on 8 ALW crystal structures together with 18 strong-binder sequences. (d) Per-position classification agreement for panel c. In both enriched settings, RACER-m shows improved concordance at flanking sites while retaining sensitivity to the $Pro^5/Asp^6$ hot spot.

#### AlphaFold3-augmented training improves global agreement with experiment

Finally, we examined the effect of AlphaFold3 structural enrichment directly across all peptide positions. Figure S4 shows: panel a, per-position correlation; panel b, per-position agreement; and panel c, the aggregate correlation when all 190 peptide mutants are pooled together. This analysis highlights that AlphaFold-enriched training substantially improves prediction consistency across positions, reducing the positional bias of the baseline model.

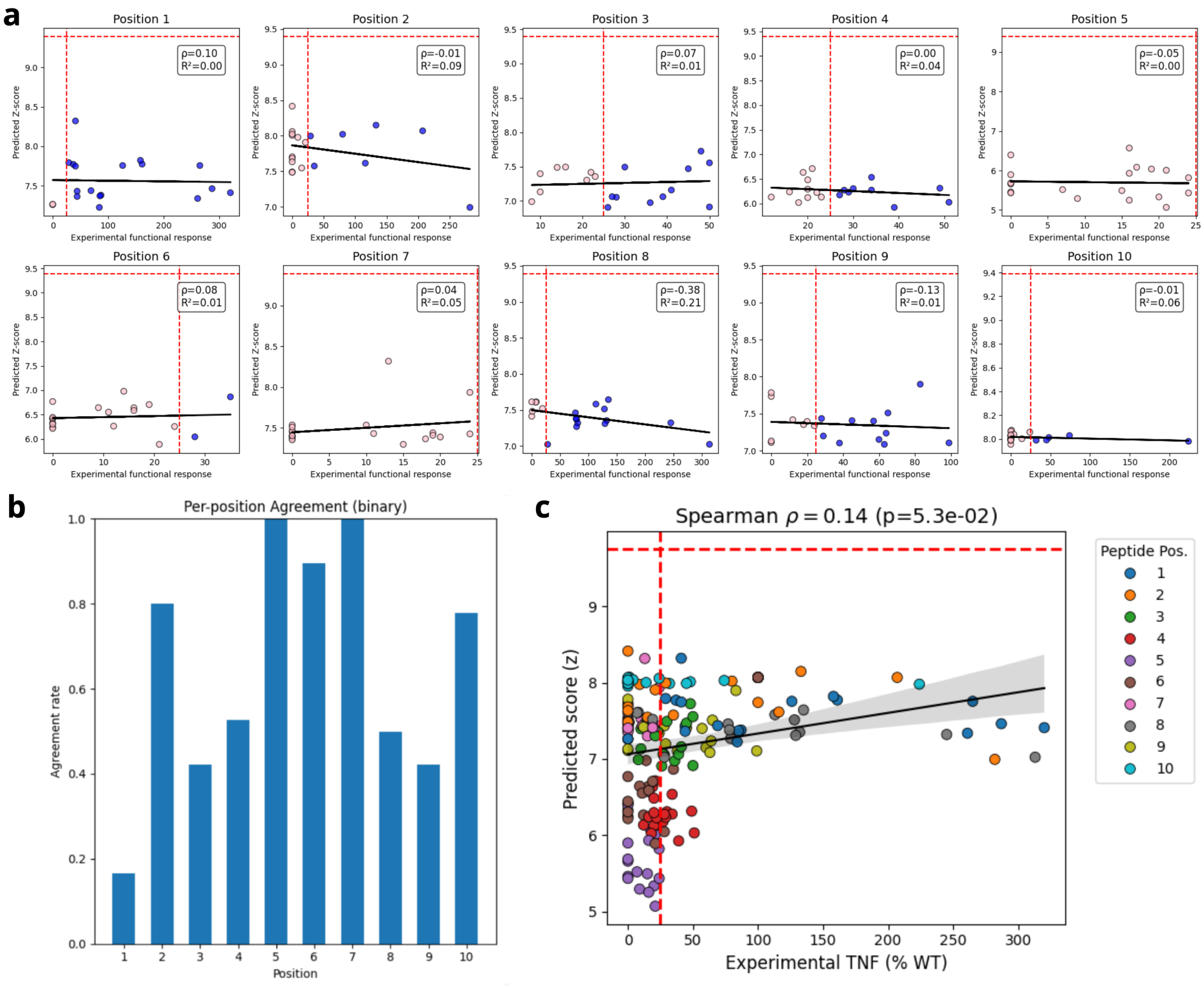


Figure S4: AlphaFold-enriched training analysis. (a) Per-position correlation. (b) Per-position classification agreement. (c) Aggregate correlation across all peptide positions.

To assess the structural relationship of the additional AlphaFold-generated models, we computed a mutual $Q$ correlation matrix across the 69 ATLAS crystal structures and the 18 AlphaFold-generated ones (Figure S5a). The clustering pattern shows that while the AlphaFold-derived 1E6–ALW structures share the closest similarity with empirical 1E6–ALW structures, they also concentrate in a distinct location in structural space relative to the crystallographic templates, indicating that they provide complementary rather than redundant coverage of the conformational landscape and are therefore useful for training augmentation. Together, panels (a–b) demonstrate that AlphaFold3-derived structures expand the structural diversity of the training set and translate this diversity into specific and localized changes in the learned interaction model. In particular, panel (b) shows that inclusion of AlphaFold3-derived 1E6–ALW structures induces systematic reweighting of the RACER-m energy matrix concentrated at peptide-facing residue pairs that directly participate in TCR recognition, while leaving the majority of non-contacting or MHC-facing interaction terms largely unchanged. These findings reinforce the role of structural augmentation in refining model sensitivity to peptide geometry rather than globally reshaping the energy landscape, thereby providing a clear mechanistic basis for the improved global agreement with experiment observed in Figure S4.

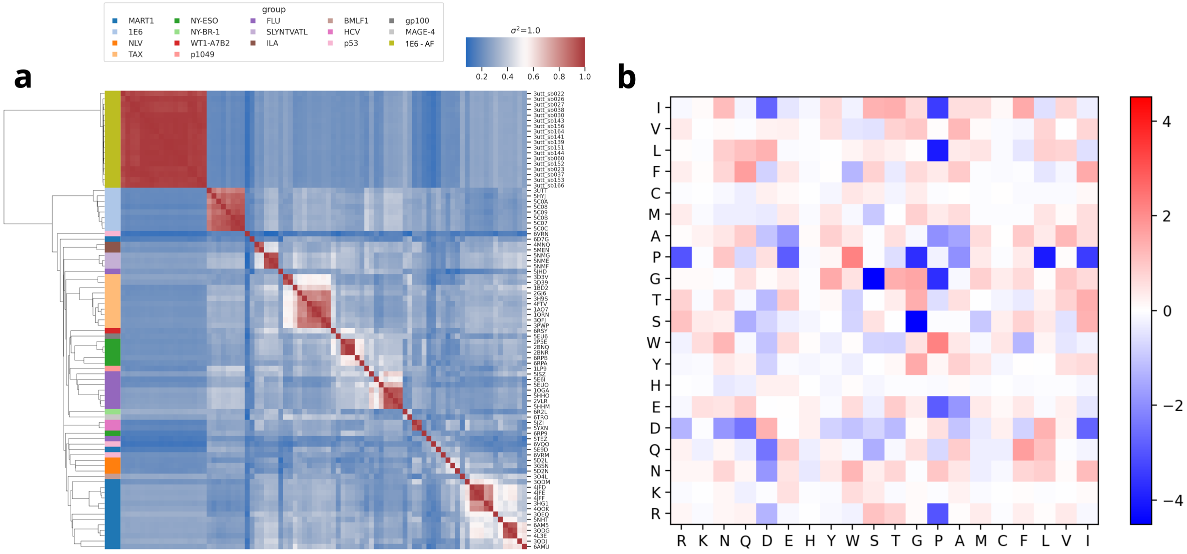


Figure S5: Mutual $Q$ correlation and energy-matrix analysis for AlphaFold3-augmented training. (a) Mutual $Q$ correlation matrix across the 69 ATLAS crystal structures together with the 18 AlphaFold3-derived 1E6–ALW structures. (b) Difference between the RACER-m energy matrices learned with (*ALW–AF18*) and without (*ALW–ATLAS8*) AlphaFold3-derived structural augmentation, highlighting localized reweighting of peptide-facing interaction terms induced by structural enrichment.

#### Dual-contact compensatory regime at the $Pro^5$ peptide hotspot

We further analyzed peptide mutations at positions where both CDR3$\alpha$ and CDR3$\beta$ directly contact the peptide, focusing on RACER-m-identified compensatory mutations that could rescue immune loss of a peptide escape variant. A representative example involves substitutions at $Pro^5$, a central hot-spot residue that anchors the peptide within the TCR interface. The compensatory complex shown in Figure S6 illustrates a predicted rescued binding configuration in which $Pro^5$ engages residues from both CDR3 loops, forming a dual-contact geometry. This architecture imposes strong structural constraints on compensatory mutational space, as both chains contribute directly to peptide recognition.

Consistent with this dual-contact geometry, RACER-m predicts that single-site edits in either CDR3$\alpha$ or CDR3$\beta$ alone are generally insufficient to restore strong binding once $Pro^5$ is perturbed. The restricted nature of compensatory solutions is quantified in the heatmap shown in Figure S7, which summarizes an exhaustive scan of compensatory substitutions across CDR3$\alpha$ positions in the presence and absence of mutations at CDR3$\beta$ position 5. When CDR3$\beta$ is held fixed, several CDR3$\alpha$ substitutions (including S$\to$Q, K$\to$Q, and S$\to$V) recover strong-binder predictions. In contrast, when CDR3$\beta$ is also mutated, compensatory solutions become rare. Among all tested combinations, only the paired substitution CDR3$\beta$ W$\to$L together with CDR3$\alpha$ S$\to$Y yields a rescued complex that re-enters the strong-binder regime.

Together, these analyses demonstrate that dual-contact hot-spot residues impose a cooperative constraint on compensatory TCR mutations, sharply limiting the space of viable rescue configurations.

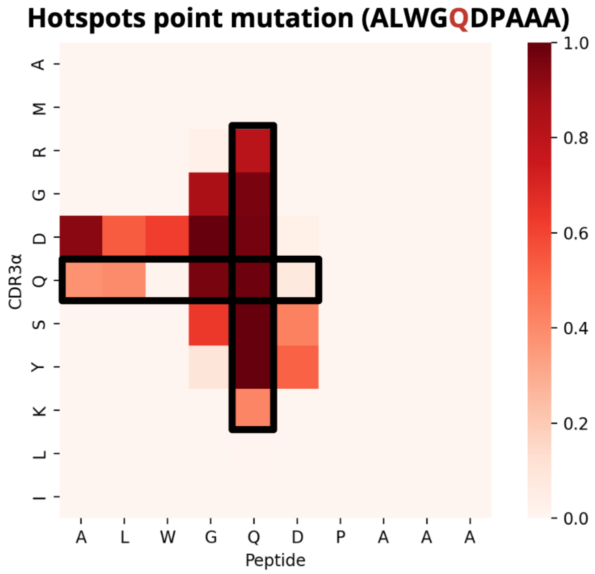


Figure S6: **Dual-contact geometry of a compensatory $Pro^5$ rescue complex.** Contact map of a representative compensatory TCR–pMHC complex in which recognition of a $Pro^5$ peptide mutant is restored. The map shows direct interactions between the peptide and both CDR3$\alpha$ and CDR3$\beta$ loops, defining a dual-contact recognition regime that constrains compensatory mutational flexibility.

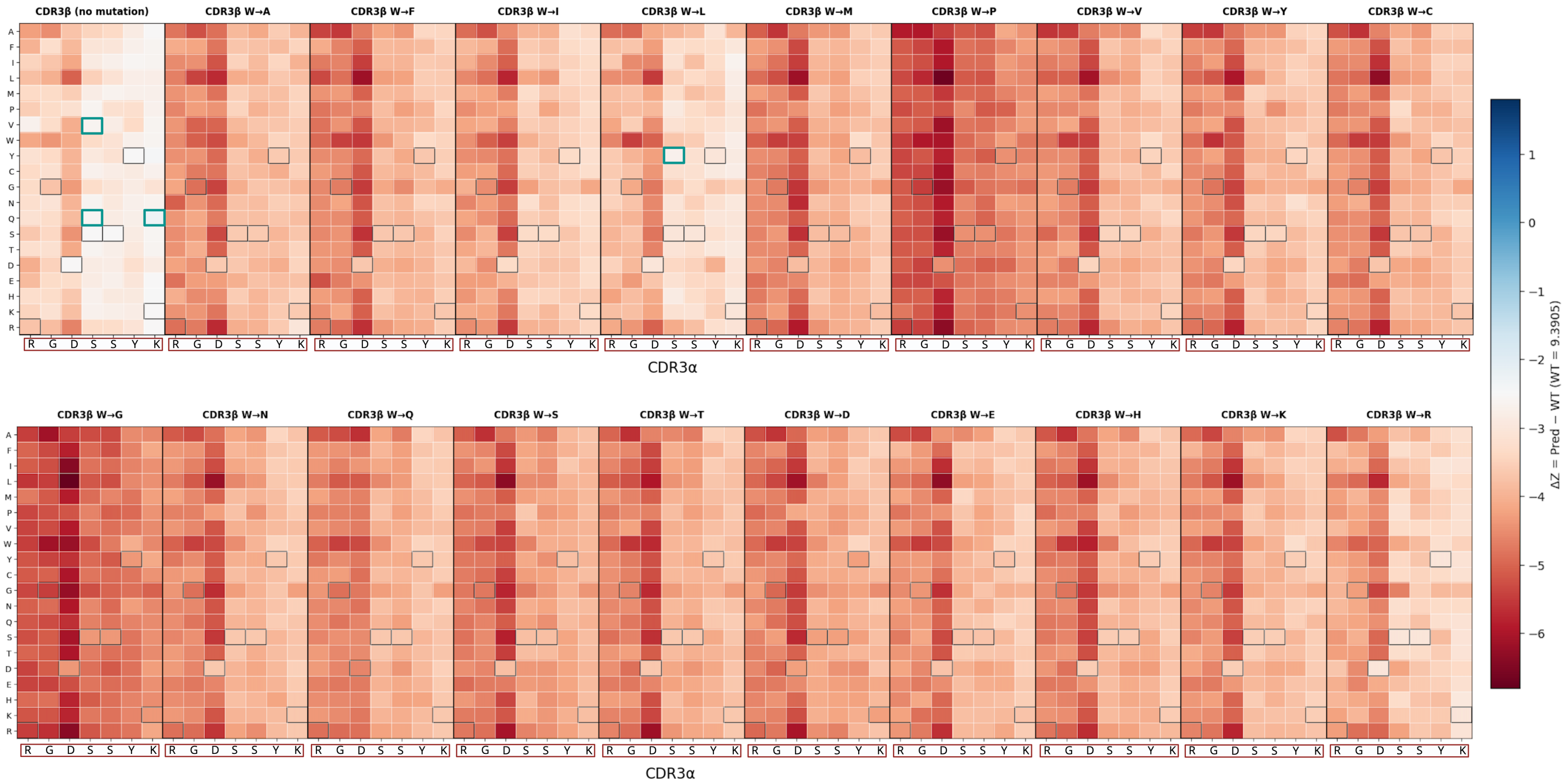


Figure S7: **Compensatory landscape leading to a dual-contact Pro$^5$ rescue.** Heatmap summarizing the exhaustive compensatory scan used to identify the rescued complex shown in Figure S6. CDR3$\beta$ position 5 was mutated, and compensatory substitutions were evaluated across the seven CDR3$\alpha$ residues contacting the peptide. Blue cells denote substitutions predicted to restore strong binding, whereas white cells indicate non-compensatory edits. The sparse distribution of successful rescues reflects the strong cooperative constraint imposed by dual-contact peptide engagement.

#### Workflow for targeted structural augmentation

The above analysis motivated a systematic workflow to determine when additional structures are needed. Toward this end, we begin with training RACER-m using empirically available antigen-matched structural data and evaluate model-predictions to empirical ground-truth for the TCR interaction with closely related (point-mutated) antigen variants. Given a set of training structures $\mathcal{D}_{\text{train}}$, we assess an in silico-derived structure obtained for the TCR interacting with peptide mutant $\tilde{s}$ based on two quantities. The first quantity, *structural coverage*, $C(\tilde{s})$, evaluates the proximity of the newly derived structure created for peptide mutant $\tilde{s}$ relative to existing training structures $s \in \mathcal{D}_{\text{train}}$ based on mutual Q $Q(\tilde{s}, s)$ structural similarity between structures $\tilde{s}$ and $s$ [3]. Specifically,

$$C(\tilde{s}) = \max_{s \in \mathcal{D}_{\text{train}}} Q(\tilde{s}, s),$$

such that a high $C(\tilde{s})$ indicates that the newly generated structure is already well represented by the existing training set, whereas a low value indicates that it occupies a less well-covered region of structural space. We emphasize that mutual $Q$ is evaluated after an *in silico* structure has been generated and is therefore used to determine whether the generated structure provides additional structural information for training, rather than to determine whether the mutant sequence itself is sufficiently different based on sequence similarity alone.

We also consider a complementary quantity, *prediction consistency*, defined as:

$$\Delta E_{\text{pred}}(\tilde{s}) = \sigma \left[ E^{(s)}_{\text{pred}}(\tilde{s}) \right]_{s \in \mathcal{D}_{\text{train}}},$$

where $E^{(s)}_{\text{pred}}(\tilde{s})$ denotes the RACER-m score predicted for peptide mutant $\tilde{s}$ when evaluated using structural template $s \in \mathcal{D}_{\text{train}}$. Thus, $\Delta E_{\text{pred}}(\tilde{s})$ quantifies the sensitivity of the predicted binding score to the choice of existing structural template; a larger value indicates greater prediction variability and therefore lower prediction consistency. Importantly, this metric is computed using the existing training structures before incorporation of the newly generated structure and does not represent a comparison between separately trained RACER-m models with and without that structure.

Structural coverage and prediction consistency provide complementary criteria for assessing whether a generated structure adds useful information to the training set. Mutants with low structural coverage and high prediction variability are prioritized most strongly for structural augmentation because they are both poorly represented by the existing structural ensemble and sensitive to template choice. Conversely, mutants with high structural coverage and low prediction variability are already well represented and are therefore less informative for augmentation. AlphaFold3-derived complexes selected through this procedure are incorporated into the antigen-restricted training set, RACER-m is retrained, and performance is evaluated on held-out peptide mutants. Figure S8 summarizes this workflow.

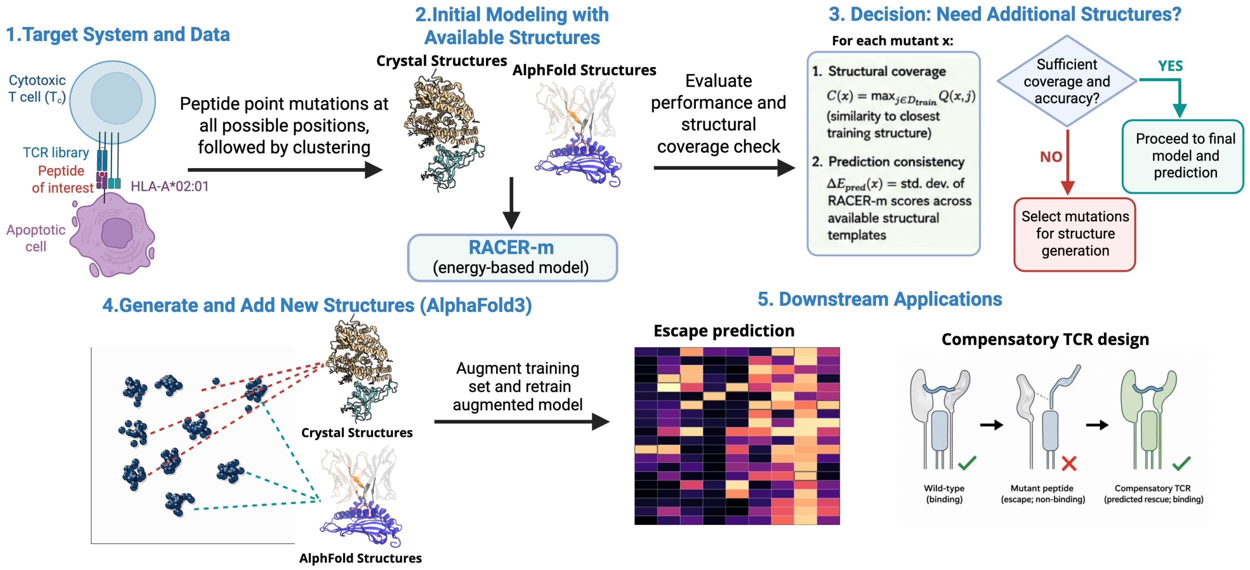


Figure S8: **Workflow for targeted structural augmentation of RACER-m.** Starting from a target TCR–pMHC system and its available mutational and structural data, RACER-m is first trained using the available antigen-matched structures. Structural coverage and prediction consistency are then assessed for each peptide mutant. When the existing structural set provides insufficient coverage or unstable predictions, representative strong-binding peptide variants are selected for AlphaFold3-based structure generation. The resulting structures are added to the antigen-restricted training set, the model is retrained, and performance is evaluated on held-out mutants.

#### AlphaFold3 validation of compensatory complexes at dual-contact hot-spot residues

To further assess the structural plausibility of compensatory solutions at dual-contact hot-spot residues, we applied AlphaFold3 to representative compensatory complexes involving substitutions at $\text{Pro}^5$. Unlike flanking or single-chain hot-spot cases, $\text{Pro}^5$ engages both CDR3$\alpha$ and CDR3$\beta$, imposing a more restrictive geometric constraint on rescue mechanisms. Figure S8 shows AlphaFold3-predicted structures for compensatory TCR edits identified by RACER-m in this dual-contact regime.

Across these cases, AlphaFold3 preserves the overall 1E6 docking geometry while reproducing the cooperative engagement of both CDR3 loops with the peptide. Compensatory substitutions predicted by RACER-m are accommodated without introducing physically implausible atomic overlaps or large-scale backbone rearrangements, but successful rescue is limited to a narrow subset of coordinated TCR edits. These structural models support the conclusion that dual-contact hot-spot residues sharply constrain compensatory flexibility and require cooperative TCR-side adaptations for recognition to be restored.

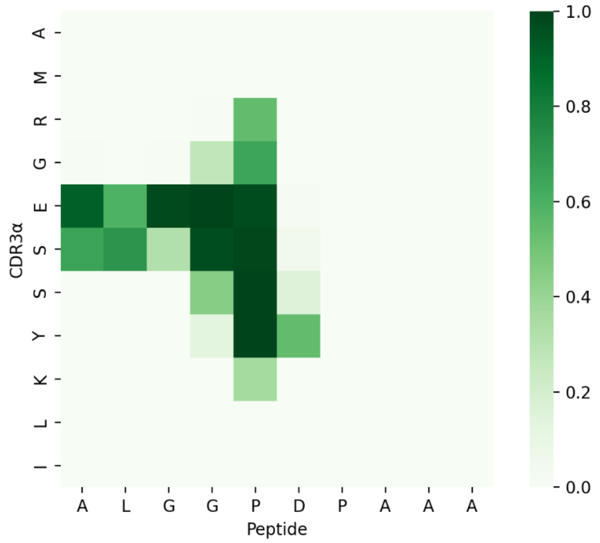


Figure S9: **AlphaFold3 structural validation of compensatory complexes at a dual-contact hot-spot residue.** AlphaFold3-predicted TCR–pMHC structures for representative compensatory solutions involving substitutions at the $\text{Pro}^5$ peptide position. The models illustrate cooperative engagement of both CDR3$\alpha$ and CDR3$\beta$ loops with the peptide, consistent with the dual-contact geometry identified by RACER-m. While the global 1E6 docking mode is preserved, successful compensatory configurations are restricted to a limited set of coordinated TCR edits, highlighting the structural constraints imposed by dual-contact hot-spot residues.

### Frustration–density distribution and peak–threshold analysis of CDR3 regions

Figure S10 provides a detailed view of how energetic frustration is distributed across the TCR interface and how this distribution changes under increasing stringency. Panels (a) and (b) quantify the number of local minima in the low-frustration contact–density profiles for the CDR3$\alpha$ and CDR3$\beta$ regions, respectively, as a function of the applied threshold. While the CDR3$\alpha$ region shows clear separation between wild-type, escape, and compensatory complexes—reflecting mutation-induced redistribution of energetic hot-spots—the corresponding curves for CDR3$\beta$ largely overlap, consistent with the conserved energetic role of this region across all variants. Panels (c–e) further illustrate these trends through residue-resolved frustration–density profiles, highlighting that compensatory mutations restore a WT-like energetic organization in CDR3$\alpha$ without introducing additional frustration elsewhere in the interface.

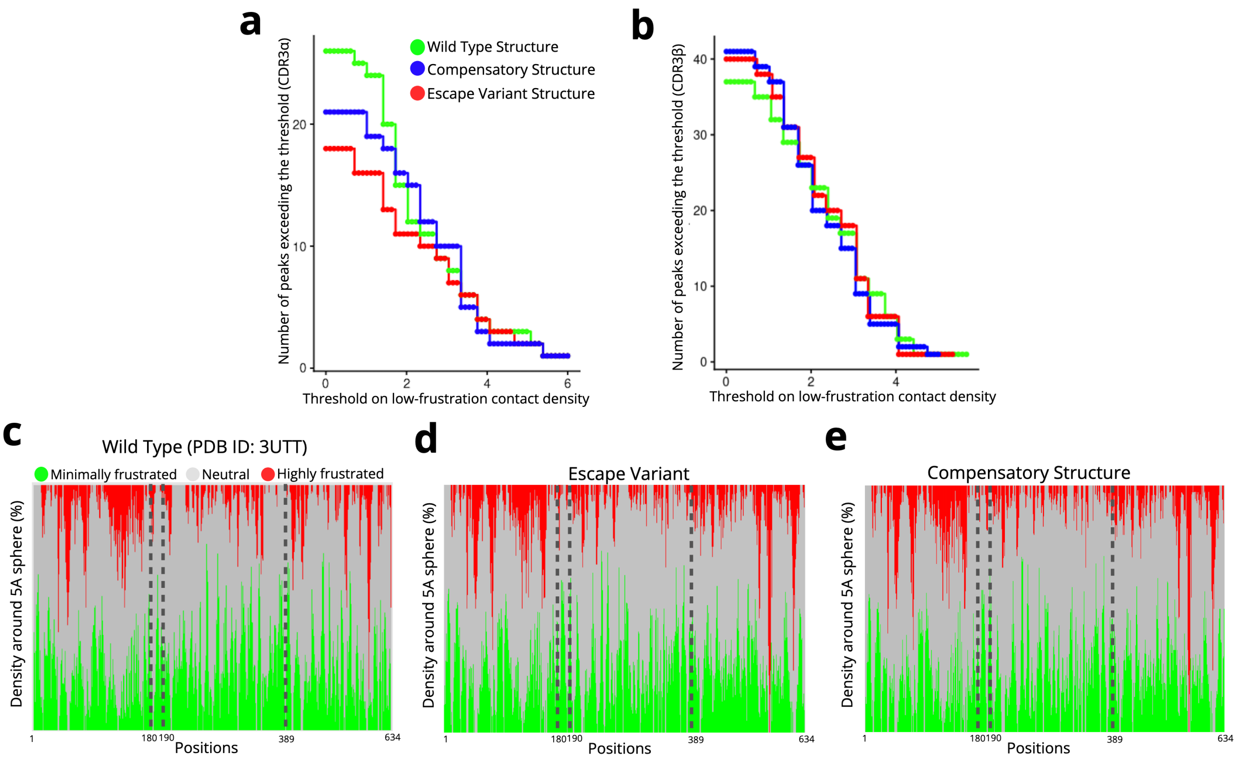


Figure S10: **Frustration–density profiles and peak–threshold analysis for CDR3 regions. (a)** Number of low-frustration peaks exceeding a given threshold within the CDR3$\alpha$ region. The number of local maxima ("peaks") in the minimally frustrated (low-frustration) contact–density profile is plotted as a function of the applied threshold for the wild-type (green), compensatory (blue), and escape (red) complexes. Each peak corresponds to a contiguous cluster of energetically favorable, minimally frustrated interactions localized along the CDR3$\alpha$ segment. **(b)** Corresponding peak–threshold analysis for the CDR3$\beta$ region, showing largely overlapping curves across all variants. **(c–e)** Residue-resolved frustration–density distributions within a 5 Å sphere for the wild-type (c), escape variant (d), and compensatory structure (e). Green, gray, and red denote minimally frustrated, neutral, and highly frustrated contacts, respectively. Gray dashed vertical lines indicate structural boundaries separating the MHC heavy chain, peptide, CDR3$\alpha$, and CDR3$\beta$ regions.

### Residue-resolved frustration density and contact maps for CDR3 regions

To further characterize the energetic organization of the TCR interface at residue-level resolution, we analyzed frustration–density profiles and contact maps for the CDR3 regions across the wild-type, escape variant, and compensatory complexes. Figure S11 focuses on the CDR3$\alpha$ region, showing both residue–residue contact maps colored by the local mutational frustration index and the corresponding frustration–density distributions within a 5 Å sphere. Consistent with the peak–threshold analysis presented in FigureS10a–b, the CDR3$\alpha$ region exhibits a clear redistribution of frustration upon introduction of the escape mutation, followed by recovery in the compensatory structure, without the emergence of new clusters of highly frustrated contacts. In contrast, the corresponding analysis for the CDR3$\beta$ region (Figure S12) reveals highly similar frustration–density profiles and contact patterns across all three complexes, indicating that the energetic landscape of CDR3$\beta$ is largely conserved and minimally affected by either escape or compensatory mutations. Finally, residue-level mutational frustration analysis at CDR3$\alpha$ position Ser94 (Figure S13) shows that substitutions to tyrosine or alanine are energetically favorable, supporting our prediction that Ser94$\mapsto$Y or Ser94$\mapsto$A can act as compensatory edits that alleviate local energetic strain and stabilize the altered interface.

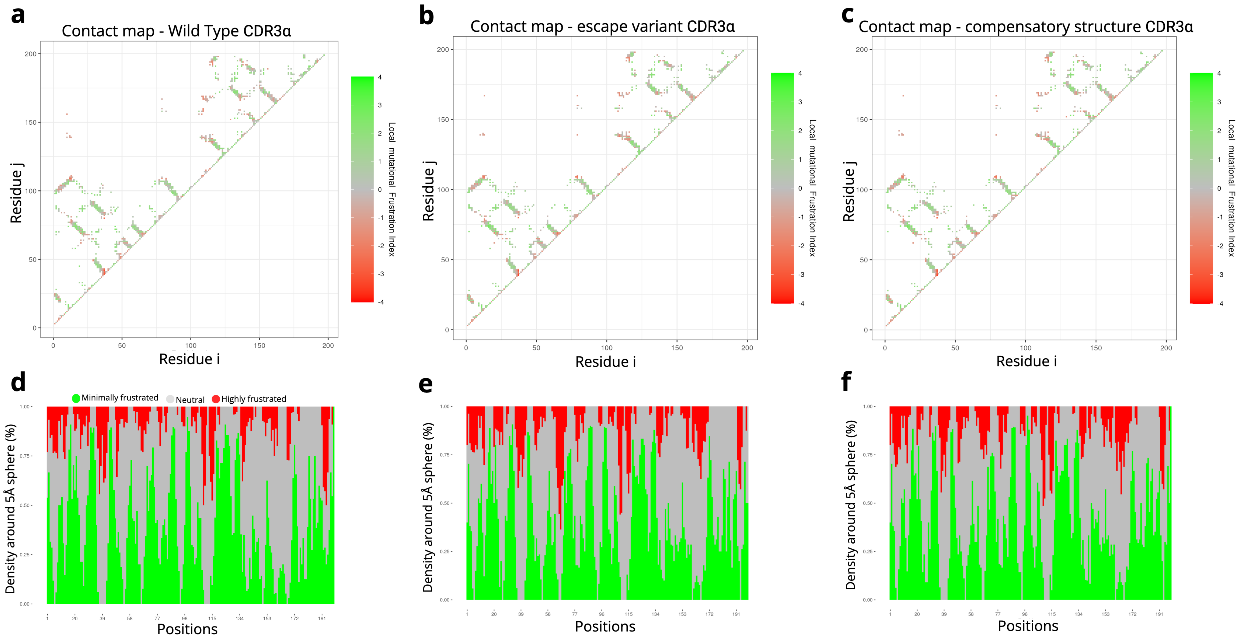


Figure S11: **Frustration density distributions and contact maps for the CDR3$\alpha$ region. (a–c)** Residue–residue contact maps colored by local mutational frustration index for the wild-type (a), escape variant (b), and compensatory (c) complexes. Green indicates minimally frustrated contacts, red indicates highly frustrated contacts, and gray denotes neutral interactions. **(d–f)** Corresponding residue-resolved frustration–density profiles computed within a 5 Å sphere around each position for the wild-type (d), escape variant (e), and compensatory (f) complexes.

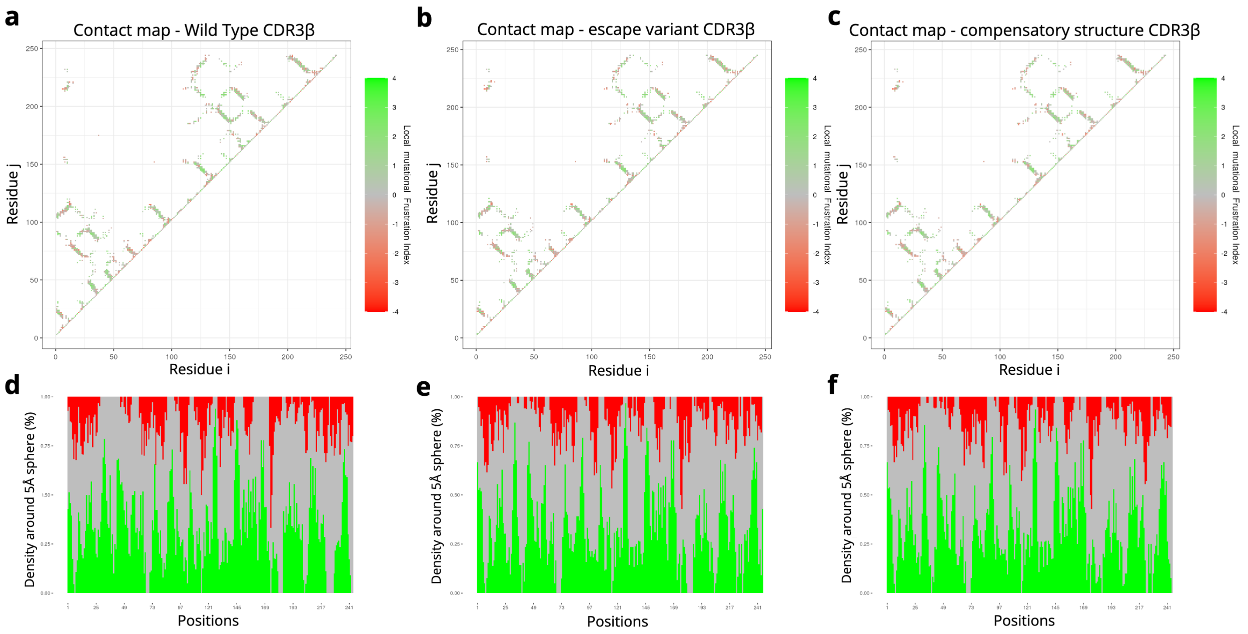


Figure S12: **Frustration density distributions and contact maps for the CDR3$\beta$ region. (a–c)** Residue–residue contact maps colored by local mutational frustration index for the wild-type (a), escape variant (b), and compensatory (c) complexes. **(d–f)** Corresponding residue-resolved frustration–density profiles within a 5 Å sphere for each complex. Across all variants, both the contact maps and frustration–density profiles for CDR3$\beta$ show strong overlap, indicating a conserved energetic landscape consistent with the limited sensitivity of this region to escape or compensatory mutations.

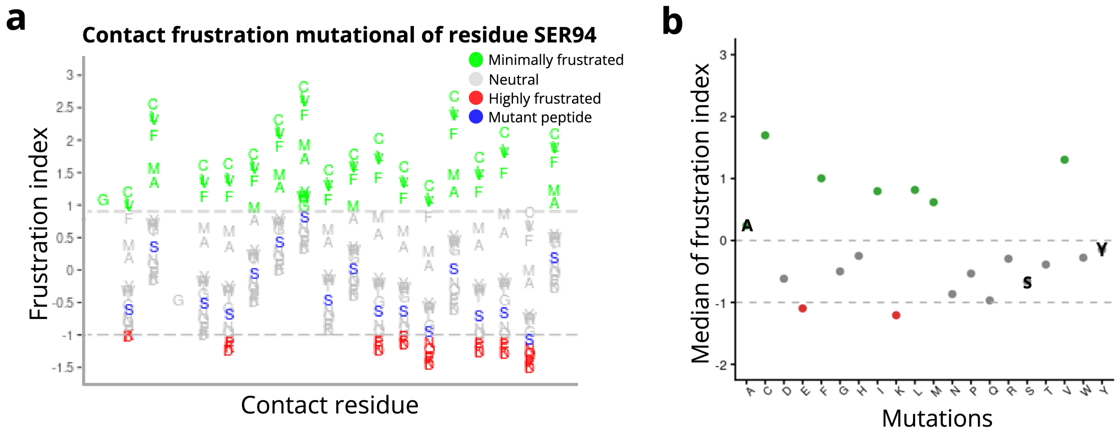


Figure S13: **Residue-level mutational frustration analysis at CDR3$\alpha$ position Ser94. (a)** Contact-level mutational frustration profiles for residue Ser94 in the CDR3$\alpha$ region. Each column corresponds to a residue contacting Ser94, and letters indicate the frustration index associated with all possible amino-acid substitutions at this position, colored by frustration class (minimally frustrated, green; neutral, gray; highly frustrated, red). The mutant peptide background is highlighted in blue. **(b)** Median frustration index across all contacts for each Ser94 substitution, summarizing the overall energetic favorability of alternative amino acids at this position. Notably, substitutions to tyrosine (Y) and alanine (A) are predicted to yield minimally frustrated interactions, consistent with our model's prediction that Ser94→Y or Ser94→A can act as compensatory mutations that alleviate local energetic strain introduced by the escape peptide.

#### Generalization of the 1E6–ALW workflow to independent CMV and SARS-CoV-2 TCR–pMHC systems

To assess whether the antigen-restricted, structurally augmented workflow developed for 1E6–ALW extends beyond a single TCR–pMHC system, we applied the same procedure to two independently characterized viral TCR–pMHC systems. Specifically, we considered TCR81-14, which recognizes the CMV-derived HLA-A*02:01-restricted epitope NLVPMVATV, and SVAR-73, which recognizes the SARS-CoV-2 spike-derived HLA-A*02:01-restricted epitope YLQPRTFLL. For each system, RACER-MAP was trained using a limited fraction of the available clone-specific mutational data together with antigen-matched structural augmentation, and performance was evaluated on held-out peptide variants.

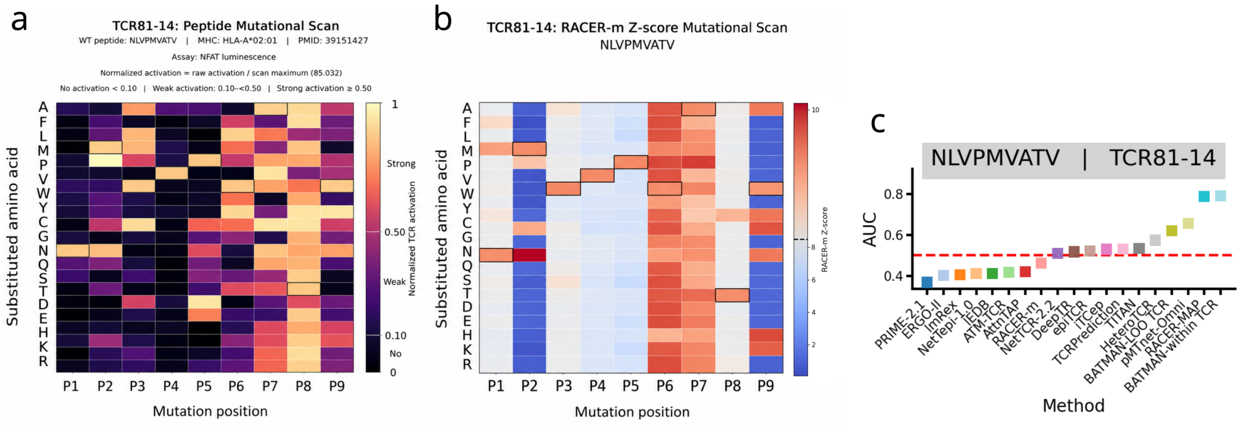


Figure S14: **Application and benchmarking of RACER-MAP for the CMV-specific TCR81-14–NLVPMVATV system.** (a) Experimental single-point mutational scan of NLVPMVATV for TCR81-14, showing normalized TCR activation across peptide positions and amino-acid substitutions. (b) RACER-MAP-predicted single-point mutational profile for TCR81-14, showing the predicted effects of amino-acid substitutions across the NLVPMVATV peptide. (c) Classification performance (AUC) of RACER-m, RACER-MAP, and published TCR–pMHC specificity predictors for distinguishing strong- and weak-activation NLVPMVATV variants. RACER-MAP was trained using only 20% of the available TCR81-14 mutational data, whereas most published comparator methods were evaluated using their existing pretrained models; BATMAN-within TCR and pTEAM used five-fold within-TCR cross-validation.

**TCR81-14 (CMV NLVPMVATV)** We first applied the workflow to the NLVPMVATV-specific TCR81-14 clone. The experimentally measured single-point mutational landscape is shown in Figure S14a, revealing the position- and substitution-dependent effects of peptide mutations on TCR81-14 activation. Using only 20% of the available TCR81-14 mutational data for antigen-specific training and structural augmentation, RACER-MAP reproduced the major features of this mutational landscape across the NLVPMVATV peptide (Figure S14b).

We next benchmarked RACER-m and RACER-MAP against published TCR–pMHC specificity predictors (Figure S14c). Most comparator methods were evaluated using their existing pretrained models and were not retrained on the TCR81-14 mutational dataset, whereas BATMAN-within TCR and pTEAM were trained and evaluated using five-fold cross-validation, corresponding to approximately 80% of the clone-specific data for training in each fold. In contrast, RACER-MAP used only 20% of the available TCR81-14 mutational data, while baseline RACER-m was trained exclusively on the ATLAS crystal-structure dataset and had no exposure to TCR81-14-specific mutational data. Despite using substantially less clone-specific training data than BATMAN-within TCR, RACER-MAP achieved comparable classification performance

(AUC = 0.786 versus AUC = 0.791 for BATMAN-within TCR), whereas baseline RACER-m performed near chance (AUC = 0.461). These results demonstrate that structural augmentation substantially improves prediction for TCR81-14 while retaining strong performance under limited clone-specific training data.

**SVAR-73 (SARS-CoV-2 YLQPRTFLL)** We next applied the same workflow to SVAR-73, a TCR clone recognizing the SARS-CoV-2 spike-derived epitope YLQPRTFLL. The experimental single-point mutational scan provides a position-resolved binding landscape across all nine peptide positions and amino-acid substitutions (Figure S15a). RACER-MAP was trained using only 20% of the available SVAR-73 mutational data together with the 163-structure ATLAS training set and AlphaFold3-derived antigen-specific structural augmentation. Importantly, the ATLAS training set contains no SARS-CoV-2 TCR–pMHC examples. The resulting RACER-MAP prediction produced a position-resolved single-point mutational landscape across all nine YLQPRTFLL positions (Figure S15).

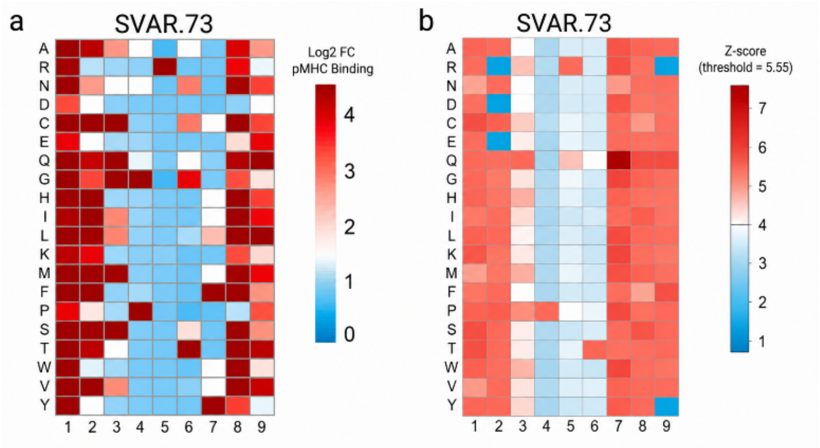


Figure S15: **Application of RACER-MAP to the SARS-CoV-2-specific SVAR-73–YLQPRTFLL system.** (a) Experimental single-point mutational binding profile for SVAR-73, showing measured TCR–pMHC recognition across all nine YLQPRTFLL peptide positions and amino-acid substitutions. (b) RACER-MAP-predicted single-point mutational profile for SVAR-73, showing predicted binding across all nine YLQPRTFLL peptide positions and amino-acid substitutions. (c) AUC comparison of RACER-MAP against eight published TCR–epitope binding predictors (AttnTAP, epiTCR, ATM-TCR, ERGO-II, ImRex, iTCep, BATMAN) for classifying strong- versus weak-binding YLQPRTFLL variants recognized by SVAR-73. RACER-MAP was trained using 20% of the available SVAR-73 mutational data together with the 163-structure ATLAS training set and antigen-specific structural augmentation, and achieved the highest AUC (0.843) among all compared methods. All benchmark methods except BATMAN were evaluated using their publicly released pretrained weights with no SVAR-73-specific training; BATMAN was retrained using only the same 20% SVAR-73 training fold, excluding its default cross-TCR hierarchical pooling across other SARS-CoV-2-specific TCRs, for a training-data-matched comparison.

To benchmark RACER-MAP against existing TCR–epitope binding predictors, we evaluated classification performance on the held-out SVAR-73 variants using area under the receiver operating characteristic curve (AUC) (Figure S15c). RACER-MAP achieved an AUC of 0.843, the highest among all compared methods, despite using only 20% of the available clone-specific mutational data for training. For comparison, published TCR–epitope predictors applied out-of-the-box (using their publicly released pretrained weights, without any SVAR-73-specific retraining) achieved substantially lower performance: iTCep (AUC = 0.620), ImRex (0.600), ERGO-II (0.553 and 0.493 for the VDJdb- and McPAS-trained versions, respectively), ATM-TCR (0.465), AttnTAP (0.456 and 0.351 for the VDJdb- and McPAS-trained versions, respectively), and epiTCR (0.395). We additionally retrained BATMAN using only the matching 20% SVAR-73 training fold, with no cross-TCR SARS-CoV-2 information available during training; under this matched, clone-restricted training condition, BATMAN achieved an AUC of 0.756. Together with the TCR81-14 results, this analysis supports the applicability of the antigen-restricted, structurally augmented RACER-MAP workflow to distinct TCRs recognizing unrelated viral epitopes under limited clone-specific training conditions, and demonstrates that RACER-MAP outperforms existing predictors even when those predictors are given comparable or greater access to clone- or virus-specific training data.
